# Loss of SORCS2 is associated with neuronal DNA double-strand breaks

**DOI:** 10.1101/2021.04.28.441600

**Authors:** Katerina O. Gospodinova, Ditte Olsen, Mathias Kaas, Susan M. Anderson, Jonathan Phillips, Rosie M. Walker, Mairead L. Bermingham, Abigail L. Payne, Panagiotis Giannopoulos, Divya Pandya, Tara L. Spires-Jones, Catherine M. Abbott, David J. Porteous, Simon Glerup, Kathryn L. Evans

## Abstract

SORCS2 is one of five proteins that constitute the Vps10p-domain receptor family. Members of this family play important roles in cellular processes linked to neuronal survival, differentiation and function. Genetic and functional studies implicate SORCS2 in cognitive function, as well as in neurodegenerative and psychiatric disorders. DNA damage and DNA repair deficits are linked to ageing and neurodegeneration, and transient neuronal DNA double-strand breaks (DSBs) also occur as a result of neuronal activity. Here, we report a novel role for SORCS2 in DSB formation. We show that SorCS2 loss is associated with elevated DSB levels in the mouse dentate gyrus and that knocking out *SORCS2* in a human neuronal cell line increased Topoisomerase IIβ-dependent DSB formation and reduced neuronal viability. Neuronal stimulation had no impact on levels of DNA breaks *in vitro*, suggesting that the observed differences may not be the result of aberrant neuronal activity in these cells. Our findings are consistent with studies linking the VPS10 receptors and DNA damage to neurodegenerative conditions.

## Introduction

SORCS2 is a member of the VPS10p-domain receptor, or sortilin, family. The family comprises five multifunctional neuronal receptors: sortilin; SORLA and SORCS1-3, which are characterised by possession of a vacuolar protein sorting (VPS) 10p domain(Hermey 2009). All family members are involved in intracellular sorting and trafficking of various neurotrophic factors, transmembrane receptors and synaptic proteins, linking them to a broad range of cellular processes, including neuronal function, differentiation and synaptic plasticity(Glerup et al 2014a).

Genetic and functional analyses implicate the VPS10p-domain receptors in cognitive functions and a wide range of neurodegenerative and psychiatric disorders. Interrogation of the GWAS catalog (https://www.ebi.ac.uk/gwas/) indicates that multiple SNPs in *SORCS2* are involved in epistatic interactions that are associated (p≤5×10^−8^) with paired helical filament tau (PHF-tau) levels (Wang et al 2020).Genetic variants in *SORCS2* are also significantly associated (p≤5×10^−8^) with alcohol withdrawal (Smith et al 2018) and risk-taking behaviour (Karlsson Linnér et al 2019). In addition, there are suggestive associations (5×10^−8^<p<1×10^−5^) with ADHD (Alemany et al 2015), anorexia nervosa (Duncan et al 2017), response to antidepressants (Fabbri et al 2018), depressive and manic episodes in bipolar disorder (Fabbri and Serretti 2016), memory performance (Greenwood et al 2019), and intelligence (Davies et al 2018). Elevated *SORCS2* levels have been detected in the brains of epileptic patients, as well as in the hippocampi of wild-type mice subjected to pentylenetetrazole (PTZ)-induced kindling, a model of epilepsy (Malik et al 2019). Meanwhile, application of PTZ-induced kindling in animals lacking *Sorcs2* increased the levels of oxidative stress and led to an exacerbated oxidative stress response in primary neurons (Malik et al 2019). Increased *SORCS2* expression has also been observed in response to application of the cortisol analogue, dexamethasone (DEXA), as well as following alcohol exposure in a human neuroblastoma cell line (Smith et al 2018). In mice, loss of *Sorcs2* has been linked to a decreased phenotypic preference for alcohol and decreased alcohol withdrawal symptoms (Olsen et al 2019), suggesting a general role of the receptor in the cellular and behavioural response to multiple stressors.

During mouse development (E15.5), *Sorcs2* is expressed in the ventral hippocampus and in tyrosine-hydroxylase-positive (TH+) neurons of the midbrain. In the adult mouse brain, *Sorcs2* is strongly expressed in hippocampal, striatal and cortical neurons (Deinhardt et al 2011; Glerup et al 2014b; Glerup et al 2016). At the cellular level, in the hippocampus SorCS2 is located at the post-synaptic density (PSD) of dendrites and within synaptic vesicles (Glerup et al 2016; Ma et al 2017). Through its interactions with the BDNF receptor tyrosine kinase, TrkB, and the pro-BDNF receptor p75^NTR^, it is implicated in the induction of NMDA-dependent long-term potentiation (LTP) and depression (LTD) in the hippocampus, respectively (Glerup et al 2016). Moreover, SorCS2 traffics TrkB to the PSD in an activity-dependent manner, thus playing a role in synaptic tagging and synaptic potentiation maintenance (Glerup et al 2016). The receptor has been also implicated in the trafficking of NMDA receptor subunits to dendritic and synaptic surfaces in medium spiny neurons of the striatum (Ma et al 2017) and in pyramidal neurons of the CA2 (Ma et al 2017; Yang et al 2020). In keeping with the above findings, *Sorcs2*^*-/-*^ mice exhibit learning and memory deficits (Glerup et al 2016) and hyperactive behaviour on exposure to novelty (Olsen et al 2021).

DNA double-strand break (DSB) formation has been previously hypothesised to be involved in learning and memory in wild-type mice via a behavioural task that involved exploration of a novel environment (Suberbielle et al 2013; Madabhushi et al 2015). Suberbielle *et al*. (2013) (Suberbielle et al 2013) reported the somewhat surprising finding of increased DSB formation in the hippocampus and parietal cortex of adult wild-type mice following exploration of a novel environment. DSBs were most abundant in the DG, an important area for learning and memory. The breaks were repaired within 24 hours leading the authors to suggest that transient break formation plays a role in chromatin remodelling and regulation of gene expression necessary for learning and memory formation. Further experiments involving direct activation of the visual cortex and the striatum via exposure to visual stimuli or optogenetic stimulation, respectively, showed that increases in neuronal activity in the absence of the behavioural paradigm were sufficient for inducing DSBs. Subsequent work by others showed that neuronal activity *in vivo* (induced via a contextual fear conditioning training paradigm) and *in vitro* also resulted in higher levels of DSBs than was seen in controls (Madabhushi et al 2015). Neuronal activity-induced DSBs were found to be located in the promoters of a subset of early-response genes and mediated by the type II topoisomerase, Topoisomerase IIβ (Topo IIβ): knockdown of Topo IIβ attenuated both DSB formation and early-response gene expression following neuronal stimulation (Madabhushi et al 2015). In keeping with these findings, *in vitro* pharmacological stimulation of neuronal activity has been shown to be associated with increased DSB formation (Suberbielle et al 2013; Madabhushi et al 2015).

Given the changes in synaptic plasticity and the altered response to novelty and to stress observed in the *Sorcs2*^*-/-*^ mice, we hypothesised that loss of the receptor may lead to alterations in the number of DNA DSBs at baseline, following exploration of a novel environment and/or following a recovery period. In keeping with previous data, we detected an increase in DSB formation in the hippocampus of wild-type mice following exploratory activity and repair of these breaks after a recovery period. Compared to wild-type mice, *Sorcs2* knock-out mice had higher levels of DSBs in the DG at baseline only. Next, we investigated whether this difference would also be observed in human neurons lacking SORCS2. We used CRISPR/Cas9 genome editing to delete the gene from Lund Human Mesencephalic (LUHMES) human neurons (Lotharius et al 2002; Scholz et al 2011). We found that neurons from *SORCS2* knock-out lines had more DNA DSBs and were characterised by decreased viability compared to wild-type lines. There was no difference in the number of breaks observed in wild-type and knock-out lines following stimulation of neuronal activity.

## Materials and Methods

### Compounds and antibodies

Primary antibodies used in this study: polyclonal sheep anti-SORCS2 (AF4238, R&D Systems), monoclonal mouse anti-γH2A.X (JBW301, Millipore) and polyclonal rabbit anti-53BP1 (NB100304, Novus Biologicals). Secondary antibodies: rabbit anti-mouse Immunoglobulins/HRP (P0260, Dako), rabbit anti-sheep Immunoglobulins/HRP (P0163, Dako), Alexa Fluor^®^ 488 donkey anti-mouse IgG (H+L) (A21202, Thermo Scientific) and Alexa Fluor^®^ 568 donkey anti-rabbit IgG (H+L) (A21207, Thermo Scientific). Etoposide was purchased from Sigma (E1383).

### Animals

Mice were housed at the animal facility at Aarhus University, in groups of up to five mice per cage with a 12-h light/12-h dark schedule and fed standard chow (1324, Altromin) and water *ad libitum*. Cages were cleaned and supplied with bedding and nesting material every week. *Sorcs2*^*−/−*^ mice had been backcrossed for ten generations into C57BL/6J Bomtac background (Glerup et al 2014b). All experiments were approved by the Danish Animal Experiments Inspectorate under the Ministry of Justice (Permits 2011/561-119, 2016-15-0201-01127 and 2017-15-0201-01192) and carried out according to the ARRIVE guidelines. Behavioural experiments were carried out using sex- and age-matched mice (male, 5-6 months old). Each of the behavioural tests described below were carried out using naïve animals in a randomized order by an investigator blinded to the mouse genotype. No animals were excluded from the subsequent analysis.

### Exploration of a novel environment

Mice in the control group (here defined as ‘home cage’) were kept in their original cages. Mice in the novel environment (‘novel environment’) and the recovery from the novel environment (‘recovery’) groups were transferred to the testing room, where they were individually exposed to a novel environment. The novel environment consisted of an Open Field Arena with four different novel objects and mint-like odour. Individual mice were allowed to explore the novel environment for 2h. After the novel environment exploration, the mice in the novel environment group were sacrificed, while the mice in the recovery group were returned to their home cages, where they recovered from the behavioural task for 24h before being sacrificed. The mice from the home cage group were sacrificed at the same time point.

### Perfusion and tissue processing

Mice were perfused transcardially with cold PBS containing heparin (10,000 U/L), followed by ice-cold 4% paraformaldehyde (PFA) in phosphate-buffered saline (PBS). Whole brains were dissected and post-fixed overnight in 4% PFA in PBS. Following post-fixation, brains were rinsed in sterile PBS and cryoprotected first in 10% sucrose and then in 30% sucrose at 4°C until the tissue sank to the bottom of the tube. Brains were subsequently embedded in OCT compound on dry ice and stored at -80°C. Coronal sections (14μm thick) containing the brain areas of interest (i.e., DG was sampled from three regions: - 1.755mm, - 2.155mm and - 2.555mm relative to Bregma; CA2 and CA3 were sampled from two regions: - 1.755mm and - 2.1550mm relative to Bregma) were obtained and mounted on Superfrost slides. Slides were stored at −80°C.

### LUHMES culture

LUHMES is a karyotypically normal human foetal mesencephalic cell line conditionally immortalised with the v-myc oncogene. Proliferation of the neuronal precursor cells can be terminated by adding tetracyclin, thus halting v-myc expression. Subsequent addition of GDNF results in robust differentiation into post-mitotic dopaminergic neurons within five days. LUHMES cells (ATCC, RRID: CVCL_B056) were grown and differentiated as described previously (Scholz et al 2011). Briefly, cell culture dishes were pre-coated with PLO (1mg/ml; P3655, Sigma) and fibronectin (1mg/ml; F1141, Sigma) in distilled H_2_O (dH_2_O) for at least 3h at 37°C. Following incubation, the coating solution was aspirated, and plates/flasks were washed two times with dH_2_O and completely air dried before cell seeding. Prior to differentiation, LUHMES cells were maintained in proliferation medium consisting of Advanced DMEM/F12 (12634028, Life Technologies), L-glutamine (200mM; 25030081, Life Technologies), N2 supplement (100x; 17502-048, Life Technologies) and b-FGF (160μg/ml; 571502, Biolegend). Experiments were conducted after 6 or 14 days of differentiation initiated by growing cells in differentiation media consisting of Advanced DMEM/F12, L-glutamine (200mM), N2 supplement (100x), cAMP (100mM; D0627, Sigma), Tetracycline hydrochloride (1mg/ml; T7660, Sigma) and recombinant human GDNF (20μg/ml; 212-GD-010, R&D). All experiments were initiated with n=9 lines for each genotype, however, occasionally the neurons “lifted” from the plastic/coverslip and that line was lost.

### CRISPR/Cas9 Genome Editing

Guide RNAs (gRNAs) targeting *SORCS2* exon 1 or exon 3 were designed using two independent online tools: the Zhang Lab CRISPR Design website (https://crispr.mit.edu) and CHOPCHOP (https://chopchop.cbu.uib.no/), and were selected based on their on/off-target activity. The oligos were phosphorylated and subsequently cloned into the px458 vector, co-expressing the Cas9 endonuclease and GFP (RRID: Addgene_48138). Low passage LUHMES cells were fed with fresh proliferating media 2h prior to transfection. Cells were dissociated using TrypLE (12605036, Thermo Scientific), counted and 2×10^6^ cells were transfected using the Basic Nucleofector kit for primary neurons (VAPI-1003, Lonza) and the D-33 programme on the Amaxa Nucleofector II B device (Amaxa Biosystems). 500μl of pre-warmed RPMI media (BE12-752F, Lonza) was added following nucleofection. The cells were then incubated at 37°C for 5min and gently added to precoated 6-well plates containing 2ml of freshly made proliferation medium. 2μg of the Cas9 plasmid containing the gRNA of interest were used in each transfection. Empty vector (EV) control lines were generated by transfecting proliferating LUHMES at an equivalent passage number with the px458 vector alone.

Forty-eight hours following transfection, cells were lifted as described before and centrifuged at 90g for 10min. The cell pellets were resuspended in 500μl of warm PBS and GFP+ cells were sorted by FACS into pre-coated 96-well plates, containing 100μl of freshly prepared proliferation medium. After seven days, 100μl of fresh proliferation medium was added to each well, and three days later single cell colonies were identified. At this stage, one third of the cells was kept for genotyping, and the rest were split into two wells of a 24-well plate for further expansion.

### CRISPR/Cas9 sgRNAs and Primer Sequences

gRNA *SORCS2* exon 1: CGGAGTGGCTTCGCGGGCGC

gRNA *SORCS2* exon 3: CCGTCATCGACAATTTCTAC

*SORCS2* exon 1 Forward primer: CCTTTCTCTGCGCTCTCG

*SORCS2* exon 1 Reverse primer: CCGCCCCTGATGACCATA

*SORCS2* exon 3 Forward primer: CAGAGTGCCCAGGACTGTAC

*SORCS2* exon 3 Reverse primer: ATGTGCCCTAGGTATGCAGG

### Western blotting

Cells were lysed in ice cold 1% Triton lysis buffer (20mM Tris-HCl pH 8.0, 10mM EDTA, 1% Triton X-100 and 1x protease inhibitor cocktail (5892970001, Roche)) and protein concentration was measured using Bio-Rad BSA protein assay (5000116, Bio-Rad). Protein lysates were loaded on NuPAGE Tris-acetate 3-8% precast gels (EA03752BOX, Life Technologies) and ran at 150V for 1.5h. Gels were transferred onto PVDF membranes at 30V for 1.5h. Membranes were blocked in 5% milk in 0.2% Tween-20 in TBS for 1h at room temperature and probed with primary antibodies against SORCS2 (1:750; AF4238, R&D Systems) and GAPDH (1:10,000; MAB374, Merck) diluted in blocking solution overnight at 4°C. After washes (3x 10min) in 0.2% Tween-20 in TBS, membranes were incubated with secondary HRP-conjugated antibodies diluted 1:10,000 in blocking solution for 1h at room temperature. After another three washes with TBS-0.2% Tween-20, blots were visualised using the Pierce ECL Plus Western Blotting Substrate (11527271, Thermo Scientific) and exposed using autoradiography film. Protein lysate obtained from HEK293 cells transfected with a plasmid overexpressing a human *SORCS2* cDNA was used as a positive control.

### Immunofluorescence staining

Slides containing brain sections were thawed at room temperature, incubated for 10min in 4% PFA in PBS and then thoroughly washed for 30min in PBS containing 100mM glycine (1042011000, EMD Millipore) followed by 30min in PBS. Heat-mediated antigen retrieval was performed by placing slides in 1x sodium citrate buffer (PHR1416, Sigma), pH 6.0, and pulse-heated for 20min in the citrate buffer in the microwave. Slides were allowed to cool for 20min inside the microwave, followed by 30min at room temperature. Slides were then washed 3 times (15min each wash) in PBS and incubated in blocking solution for 1.5h at room temperature. Blocking solution contained 5% normal donkey serum (D9663, Sigma), 1% BSA (421501J, VWR), 0.1% Triton-X and 0.05% Tween-20 in PBS. Slices were incubated with monoclonal mouse anti-γH2A.X primary antibody (1:50; JBW301, Millipore) in 5% normal donkey serum and 1% BSA in PBS at 4°C overnight. On the following day, slides were further incubated for 30min at 37°C and washed 3 times in PBS (15min each wash). Slides were then incubated with 3% Sudan black solution in 70% ethanol for 10min at room temperature. After 3 rinses in dH_2_O, slides were incubated with corresponding Alexa-conjugated secondary antibody (1:500; A21202, Thermo Fisher) diluted in 5% normal donkey serum in PBS for 1h at 37°C. Slides were then washed 3 times in PBS, followed by 3 times in dH_2_O (15min each wash). DAPI (D9542, Sigma) diluted 1:1,000 in PBS was subsequently applied for 10min and washed off with PBS (3 washes, 5min each). Sections were mounted in ProLong Gold antifade mountant (P36930, Thermo Scientific).

### Immunocytochemistry

Pre-differentiated (day 2) LUHMES were plated down (0.15×10^6^ cells per well) and grown on acid-etched coverslips, placed in 24-well plates and coated with PLO and fibronectin, followed by Geltrex (A1413201, Thermo Scientific). Day 14 LUHMES neurons were fixed with 4% PFA for 15min, rinsed with PBS and stored in TBS at 4°C until required. Neurons were permeabilised in 0.1% TBS-Triton X for 5min. Following three rinses with TBS, coverslips were incubated in blocking solution (5% normal donkey serum in 0.1% TBS-Tween) for 1h at room temperature and then overnight at 4°C with primary antibodies diluted in blocking solution. The next day, neurons were washed with 0.1% Tween-TBS (3×10min) and incubated with corresponding secondary antibodies for 1h at room temperature. Secondary antibodies were diluted, together with DAPI (1:1,000; D9542, Sigma), in 4% normal donkey serum in 0.1% TBS-Tween. Cells were washed with TBS (3×10min) and mounted with ProLong Gold antifade mountant (P10144, Thermo Scientific). Primary antibodies used in this study were: mouse monoclonal anti-γH2A.X (1:400; JBW301, Millipore), rabbit polyclonal anti-53BP1 (1:1000; NB100304, Novus Biologicals), mouse monoclonal anti-PSD93 (1:500; NBP2-58558, Novus Biologicals), mouse monoclonal anti-synaptophysin (1:500; SMC-178D, StressMarq Bio.) and rabbit monoclonal anti-GluR1 (1:500; 04-855, Millipore). Secondary antibodies were Alexa Fluor 488-donkey anti-mouse IgG (1:300; A21202, Thermo Scientific), Alexa Fluor 596-donkey anti-rabbit IgG (1:500; A21207, Thermo Scientific) and Alexa Fluor 647 Phalloidin (1:1000; A22287, Thermo Scientific).

### Treatments

For the etoposide treatment experiments, pre-differentiated (day 2) wild-type and *SORCS2* knock-out LUHMES were plated down (0.15×10^6^ cells per well) and differentiated until day 14. LUHMES neurons were incubated with 0.5μM etoposide (E1383, Sigma) for 4h at 37°C prior to fixation. For the experiments involving stimulation with glycine, pre-differentiated (day 2) wild-type and *SORCS2* knock-out LUHMES were plated down (0.05×10^6^ cells per well) and differentiated until day 14. LUHMES neurons were incubated in a Mg^2+^ - free ACSF (125mM NaCl, 2.5mM KCl, 26.2mM NaHCO_3_, 1mM NaH_2_PO_4_, 11mM glucose, and 2.5mM CaCl_2_) supplemented with 300µM glycine (Sigma Aldrich) for 5 min, followed by a 15 min incubation in ACSF containing 1.25mM MgCl_2_ at 37°C prior to fixation.

### Image acquisition and analysis

All imaging and counting procedures were performed blind to genotype. Image analysis was performed using the software package Fiji. Z-stacked confocal images, with a step size of 0.25µm (brain sections) or 1μm (LUHMES neurons), were acquired on a Nikon STORM/A1+ microscope at 60x (brain sections) or 100x (LUHMES neurons) magnification, using the NIS Elements software. The optimal laser intensity and gain that gave no signal in the no-primary antibody controls, were established and kept constant for all subsequent analyses. Three images of each region of interest were obtained from each mouse. The number of neurons with one or more γH2A.X-positive foci, as well as the total number of nuclei within a given area (approximately 200 nuclei on average) were counted manually and the percentage of γ-H2A.X-positive nuclei determined for each image. In the case of LUHMES neurons, nine independent wild-type and nine independent *SORCS2* knock-out lines were analysed. Approximately 100 nuclei (from four images belonging to different regions of the same coverslip) were counted for each line, and the number of γH2A.X/53BP1-positive foci per nucleus was calculated.

### Quantitative reverse transcriptase PCR (qRT-PCR)

Cell pellets from day 14 LUHMES neurons were resuspended in RLT buffer (Qiagen) with 10% (v/v) 2-mercaptoethanol. Total RNA was extracted using the RNeasy mini kit (Qiagen), and 1μg per sample was reverse transcribed with Multiscribe Reverse Transcriptase using random hexamers in a 80μl reaction. Controls, in which 25ng RNA of each sample was used to make cDNA in the absence of the Multiscribe Reverse Transcriptase, were included to detect genomic contamination.

PCR amplification of the cDNA obtained for each sample was quantified using the TaqMan^®^ Universal PCR Mix No AmpErase^®^ UNG (Life Technologies), and the threshold cycle (Ct) was determined using the Applied Biosystems 7900HT Fast Real-Time PCR System and the corresponding SDS software. TaqMan probes were used for the detection of *TOP2B* and eight reference genes (*CYC1, ERCC6, SDHA, TOP1, RPLPO, SCLY, TBP and UBE4A*). The GeNorm software was used to identify the most stably expressed reference genes (*SDHA and UBE4A*). A standard curve, generated from a dilution series, was run for *TOP2B* and the reference genes. The baseline and Ct values were determined for each gene and expression levels were calculated using the standard curve method for absolute quantification, where unknowns are compared to the generated standard curve and values are extrapolated. *TOP2B* expression values were subsequently normalised to the geometric mean of the reference genes.

### Viability assay

Neuronal viability was assessed using the Alamar Blue assay (DAL1025, Thermo Scientific). This assay was chosen as: 1) it does not interfere with cell functioning and 2) it is not an end-point assay, i.e. it allows viability to be measured at multiple time points (Rampersad 2012). Viability was measured at day 6 and day 14 from an equivalent number of neurons (0.25×10^6^) per line by replacing the medium with freshly made differentiation medium containing 10% (v/v) Alamar Blue solution. Cells were incubated with the Alamar Blue solution for 2h, after which the solution was transferred to a new 24-well plate and fluorescence measured in a FLUOstar OMEGA plate reader using an excitation wavelength of 540-570nm, and an emission wavelength of 580-610nm.

### Statistical analysis

Normal distribution and variance homogeneity were assessed for each dataset (Suppl. Table 1) using the Shapiro-Wilk normality and an F test, respectively. Where linear regression models were used, normal distribution and variance homogeneity of the residuals were assessed using the Shapiro-Wilk normality test and the Spearman’s rank correlation test for heteroscedasticity, respectively. When the assumptions of normal distribution and homogeneity of variance were met, parametric tests were performed, and the data was expressed as mean ± SD. Otherwise, the data was reported as median with interquartile range and analysed using non-parametric tests. Differences between two means were assessed using unpaired Student’s t-test (two-tailed; for parametric data) or Mann Whitney test (two-tailed; for non-parametric data). Two-way ANOVA was performed when multiple means were compared. Statistical analyses were performed using GraphPad Prism. Sample sizes were estimated based on previously reported findings (Suberbielle et al 2013) or pilot experiments and calculated using the G-power software. Null hypotheses were rejected when p<0.05. Inclusion criteria were: number of animals available for the mice; number of cell lines available following genome editing and production of neurons. There were no exclusion criteria. Outlier removal was not performed.

## Results

Our goals were to investigate whether exploration of a novel environment led to a temporary increase in the number of DSBs detected in the mouse brain in our hands and ii) whether deletion of *Sorcs2* in mice leads to higher levels of DSB formation upon exploration of a novel environment and/or a deficit in break repair. The novel environment paradigm comprised three groups of mice (5-6 months of age): those that a) remained in their home cage (baseline group); b) explored a novel environment (novel environment group) and c) explored a novel environment, followed by a recovery period in the home cage (recovery group), before they were sacrificed (Fig. 1a). As described previously (Suberbielle et al 2013), the proportion of neurons positive for γH2A.X (a widely used marker of DNA DSBs in neurons and other cell types) was determined in three brain regions (DG, CA2 and CA3 of the hippocampus, Suppl. Fig. 2; Suppl. Table 2).

**Fig. 1.**
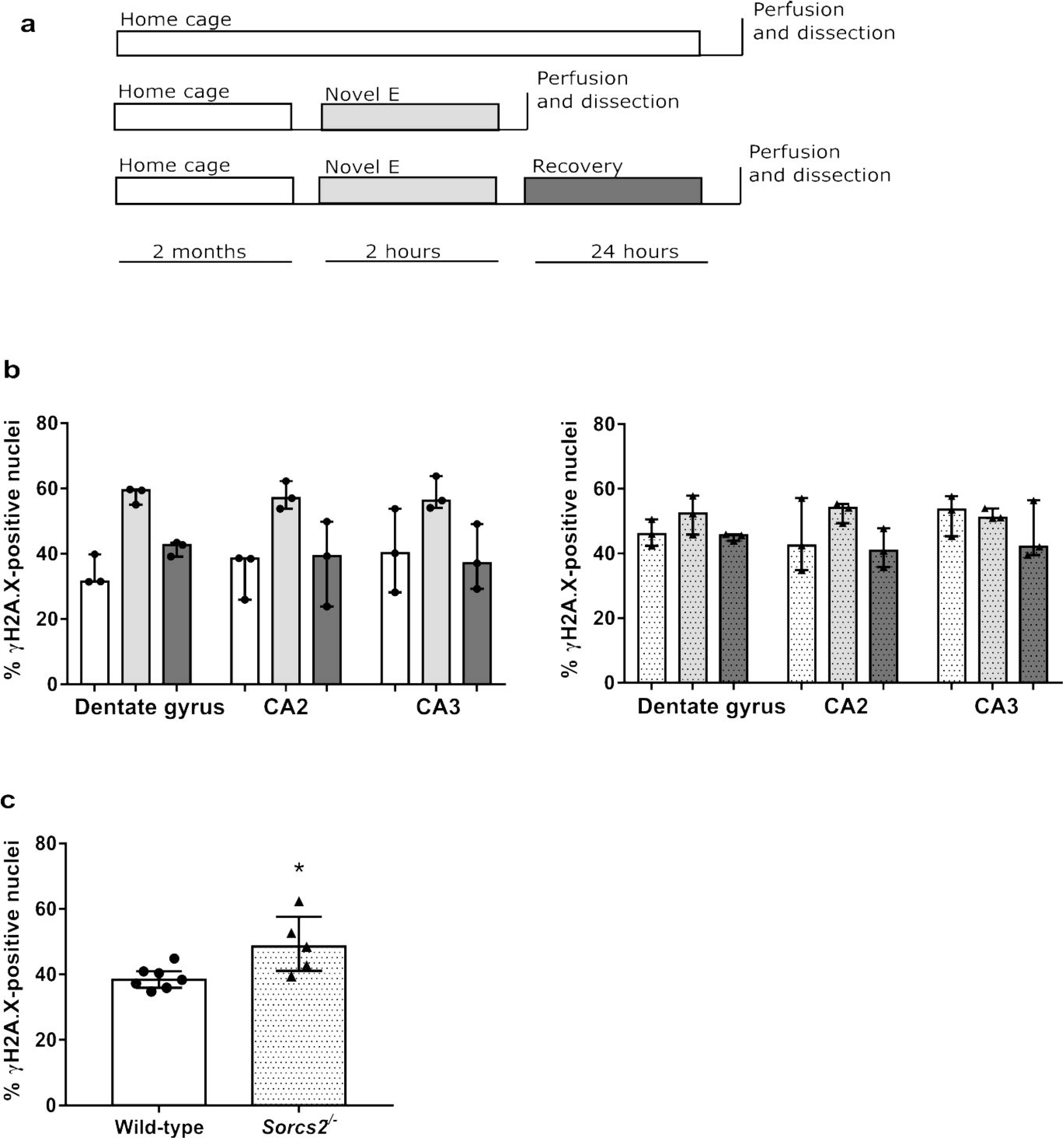
Exploration of a novel environment is associated with a transient increase in DSBs in the dentate gyrus and the CA2. (a) Experimental design. Wild-type (WT) and *Sorcs2*^*-/-*^ mice were divided into three groups: ‘home cage’ (white), ‘Novel E’ (novel environment; light grey) and ‘recovery’ (dark grey). For each brain region, the percentages of γH2A.X-positive nuclei was calculated in 5-6 month-old WT (open bars) (b) and *Sorcs2*^*-/-*^ mice (dotted bars) (c) belonging to one of the three experimental groups. Three brain sections per region per mouse, n=3 per experimental group. (d) Percentage of nuclei positive for γH2A.X in the DG of an independent set of wild-type (open bars) and *Sorcs2*^*-/-*^ (dotted bars) mice. Three brain sections per region per mouse, n=7-5. *p<0.05 (Mann-Whitney test). Error bars represent median with interquartile range

In wild-type mice in each of the three brain regions, we observed a similar pattern to that described by Suberbielle et al. (2013), i.e. wild-type mice exposed to the NE had more cells with DSBs than the mice in the baseline and the recovery groups (Fig. 1b, left). In contrast, this pattern was not present in the *Sorcs2*^*-/-*^ mice, which, to our surprise, appeared to have a greater percentage of DSB-positive nuclei at baseline (Fig. 1b, right). Given these results, we next sought to test the finding of a higher number of breaks at baseline in the DG of the *Sorcs2*^*-/-*^ mice using an independent set of age and sex-matched wild-type and knock-out ^-^ mice. We detected significantly higher levels of DSBs in the *Sorcs2*^*-/-*^ mice (U = 4, p = 0.03; Fig. 1c).

Having determined that the *Sorcs2*^*-/-*^ mice had higher levels of DNA DSBs at baseline we set out to investigate whether this phenotype was also present in human neurons lacking *SORCS2*. We used CRISPR/Cas9 genome editing (Fig. 2a) to delete the gene in the human neuronal cell line, LUHMES, a karyotypically normal foetal mesencephalic cell line that can be robustly differentiated into post-mitotic dopaminergic neurons (Suppl. Fig. 3), with the majority of cells generating trains of spontaneous action potentials after 10-12 days of differentiation (Scholz et al 2011). Loss of *SORCS2* expression was shown by western blotting (Fig. 2b; Suppl. Fig. 4). Nine independent lines were generated using two different gRNAs (four produced using a gRNA targeting exon 1 and five from the exon 3 gRNA) were used in all subsequent analyses.

**Fig. 2.**
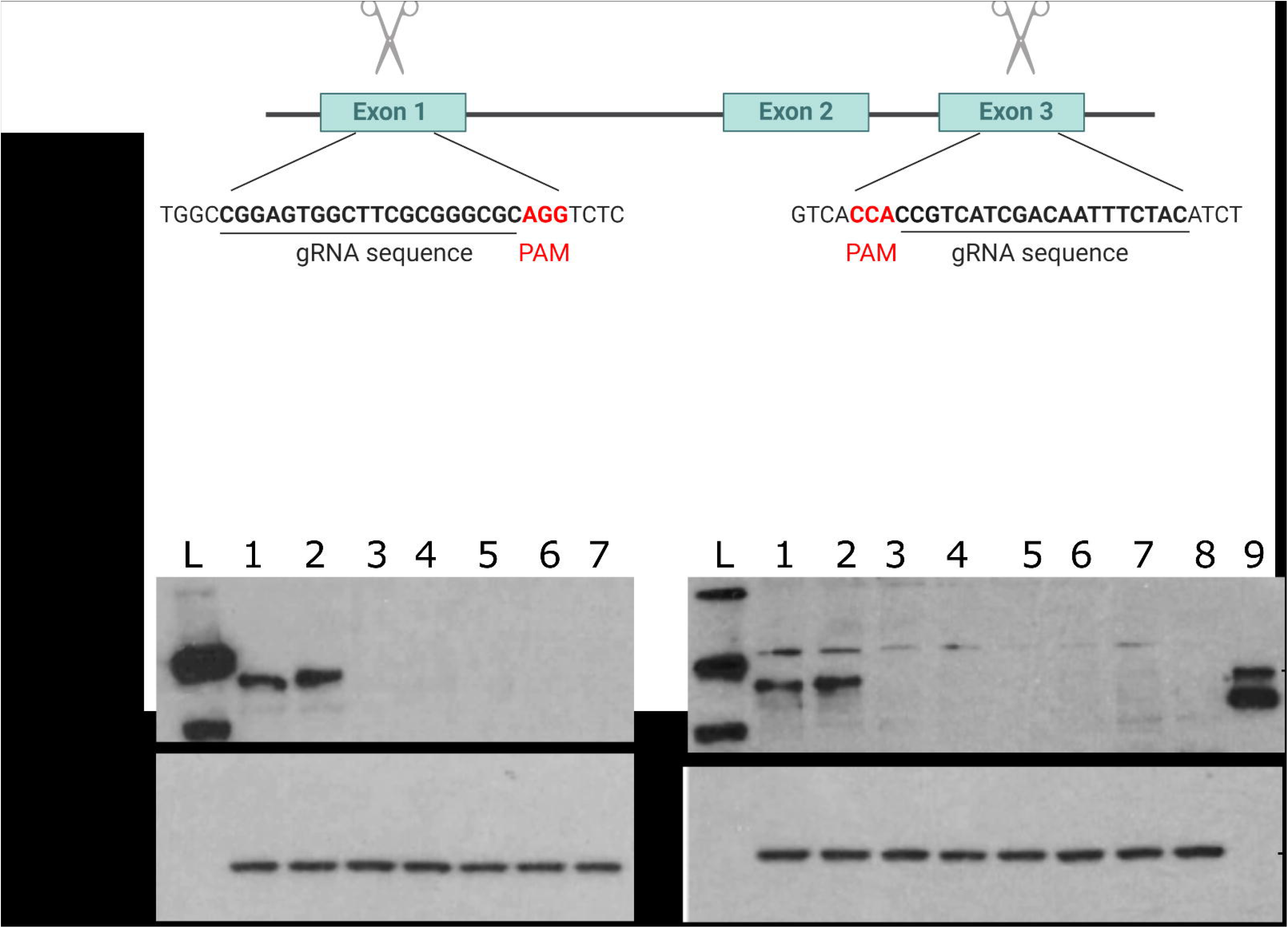
Strategy for knocking out *SORCS2 in* LUHMES using CRISPR/Cas9 genome editing. (a) Experimental design of the CRISPR/Cas9 experiments. gRNA sequences (underlined) within *SORCS2* exon 1 and exon 3 used (separately) to knock out the gene using CRISPR/Cas9 genome editing. Created with BioRender.com. (b) Representative western blots show a complete loss of SORCS2 in the knock-out (KO) clones after targeting exon 1 or exon 3. Samples loaded on the blot on the left correspond to: 1 and 2 lysates obtained from wild-type (WT) LUHMES neurons (day 14), 3-7-from *SORCS2* KO exon 1 clones 1-5 (day 14) generated by targeting exon 1. Samples loaded on the blot on the right correspond to: 1 and 2 lysates obtained from WT LUHMES neurons (day 14), samples 3-8-from *SORCS2* KO exon 3 clones 1-6 (day 14) generated by targeting exon 3. Sample 9 constitutes a positive control (protein lysate from HEK293 cells overexpressing *SORCS2*). ‘L’ stands for ladder in both blots. *SORCS2* exon 1 clone 4 and *SORCS2* exon 3 clone 5 did not survive neuronal differentiation and were not included in any subsequent experiments.

To evaluate the effect of knocking out *SORCS2* on DNA DSB formation in human neurons, we stained untreated control (consisting of wild-type (WT) and empty vector (EV) lines) and *SORCS2* knock-out LUHMES neurons (day 14) for γH2A.X and 53BP1. The latter protein is quickly recruited to DSB sites, where it binds to γH2A.X and acts as a scaffold for the binding of additional DNA repair proteins from the non-homologous end joining (NHEJ) pathway, the main DNA repair pathway in post-mitotic cells (Firsanov et al 2011). As previously reported for neurons (Crowe et al 2006), more than 90% of the analysed neurons (wild-type and knock-out) had fewer than three double positive foci per nucleus, with the majority of nuclei having no foci (Figure 3A; top row). There was no significant difference in the number of foci per nucleus between control and *SORCS2* knock-out neurons (U = 29, p = 0.34; Fig. 3b). Comparable levels of DSBs were observed between the WT and EV lines (Suppl. Fig. 5a), as well as between the *SORCS2* KOs generated by targeting exon 1 and exon 3 (U = 7, p = 0.556, Suppl. Fig. 5b). As DNA DSBs are rare, due to their dynamic repair, we next assessed whether *SORCS2* loss would have an effect on the number of DSBs following treatment with etoposide, which causes accumulation of Topoisomerase II (TopoII)-dependent DNA DSBs, by preventing their re-ligation through stabilisation of the TopoII-DNA cleavable complex (Montecucco et al 2015). As expected, etoposide treatment greatly increased the number of DSBs per nucleus in both wild-type and *SORCS2* knock-out LUHMES neurons Fig. 3a). However, comparing the number of γH2A.X/53BP1-positive foci per nucleus between etoposide-treated wild-type and *SORCS2* knock-out lines showed a significant increase in the SORCS2^-/-^ lines (t_2_= 2.148, p = 0.047;Fig. 3c). There was no significant difference in the number of γH2A.X/53BP1-positive foci per nucleus between the *SORCS2* knock-out clones derived by targeting exon 1 and those generated by disrupting exon 3 (U = 9, p = 0.905; Fig. 3d). No difference was observed between the two control groups, either, Suppl. Fig. 5c).

**Fig. 3.**
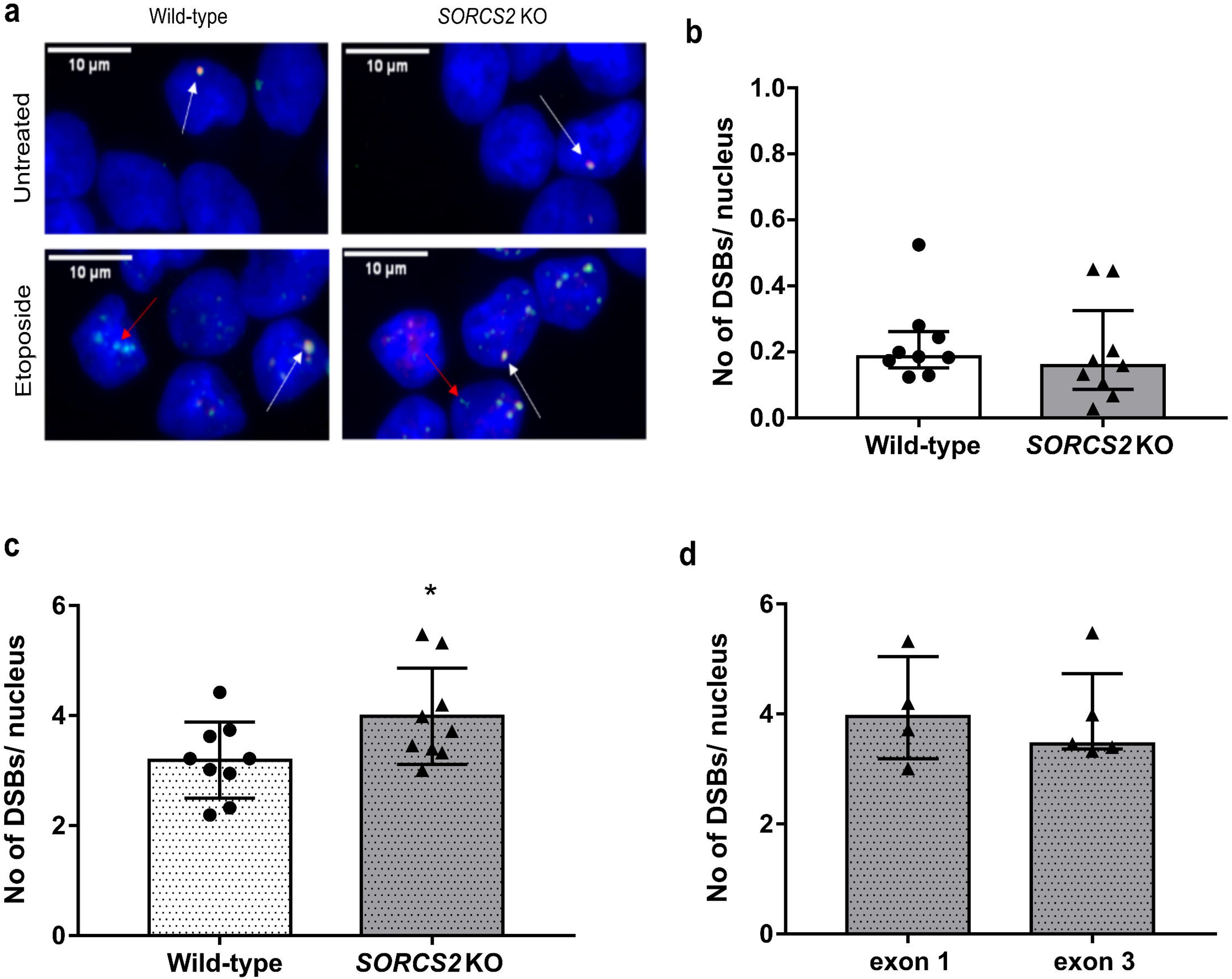
Knocking out *SORCS2* leads to increased TopoIIβ-dependent DSB formation in LUHMES neurons. (a) Representative confocal images from untreated (top row) and etoposide-treated (bottom row) wild-type (WT) and *SORCS2* knock-out (KO) LUHMES neurons (day 14) immunostained with γH2A.X (green) and 53BP1 (red), and counterstained with DAPI (blue). White arrows point towards γH2A.X/53BP1 dual positive foci, and red-towards foci positive for γH2A.X only. Images were taken at 100x magnification; scale bars: 10 μm. (b) Number of DSBs (γH2A.X/53BP1-positive foci) per nucleus in untreated WT (white bar) and *SORCS2* KO (grey bar) LUHMES neurons (day 14); n= 9 independent cell lines per genotype. Mann-Whitney test, p > 0.05. Error bars represent median with interquartile range. (c) Number of DSBs (γH2A.X/53BP1-positive foci) per nucleus in etoposide-treated (dotted bars) WT (white bars) and *SORCS2* KO (grey bars) LUHMES neurons (day 14); n=9 independent lines per genotype. * p < 0.05, Unpaired Student’s t-test; error bars represent means ± SD. (d) Number of DSBs (γH2A.X/53BP1-positive foci) per nucleus in etoposide-treated *SORCS2* KO LUHMES neurons (day 14) generated by targeting exon 1 (n=4 independent cell lines) or exon 3 (n=5 independent cell lines). Mann-Whitney test, p > 0.05; error bars represent median with interquartile range. Approximately 100 nuclei counted per cell line

Topoisomerase IIβ (TopoIIβ) is the active form of topoisomerase in terminally differentiated cells, such as neurons. Treatment with etoposide had no impact on expression levels of *TOP2B*, which encodes TopoIIβ (F_1, 16_ = 0.978, p = 0.337, Suppl. Fig. 6). In addition, there was no significant difference in *TOP2B* levels between genotypes either prior to or following etoposide treatment (F_1, 16_ = 2.652, p = 0.123, Suppl. Fig. 6).

Given the link between neuronal activity and TopoIIβ-mediated DNA DSBs (Madabhushi et al 2015), we next investigated whether an established paradigm of neuronal stimulation would have a differential impact on the formation of DNA DSBs in *SORCS2* knock-out and wild-type LUHMES neurons. Incubation with glycine (300µM) led to an increase in the number of DNA breaks (Fig. 7a and b). Treatment with glycine was also associated with a significant reduction in the somatic surface area occupied by AMPA receptors, indicative of stimulation of neuronal activity (t_7_ = 2.606, p = 0.035, Suppl. Fig. 7c and d). We next compared the impact of glycine treatment in control (WT and EV) and *SORCS2*^*-/-*^ lines. No significant difference in the number of DNA DSB foci was observed between the two groups (t_14_ = 0.383, p = 0.708, Fig. 4).

**Fig. 4.**
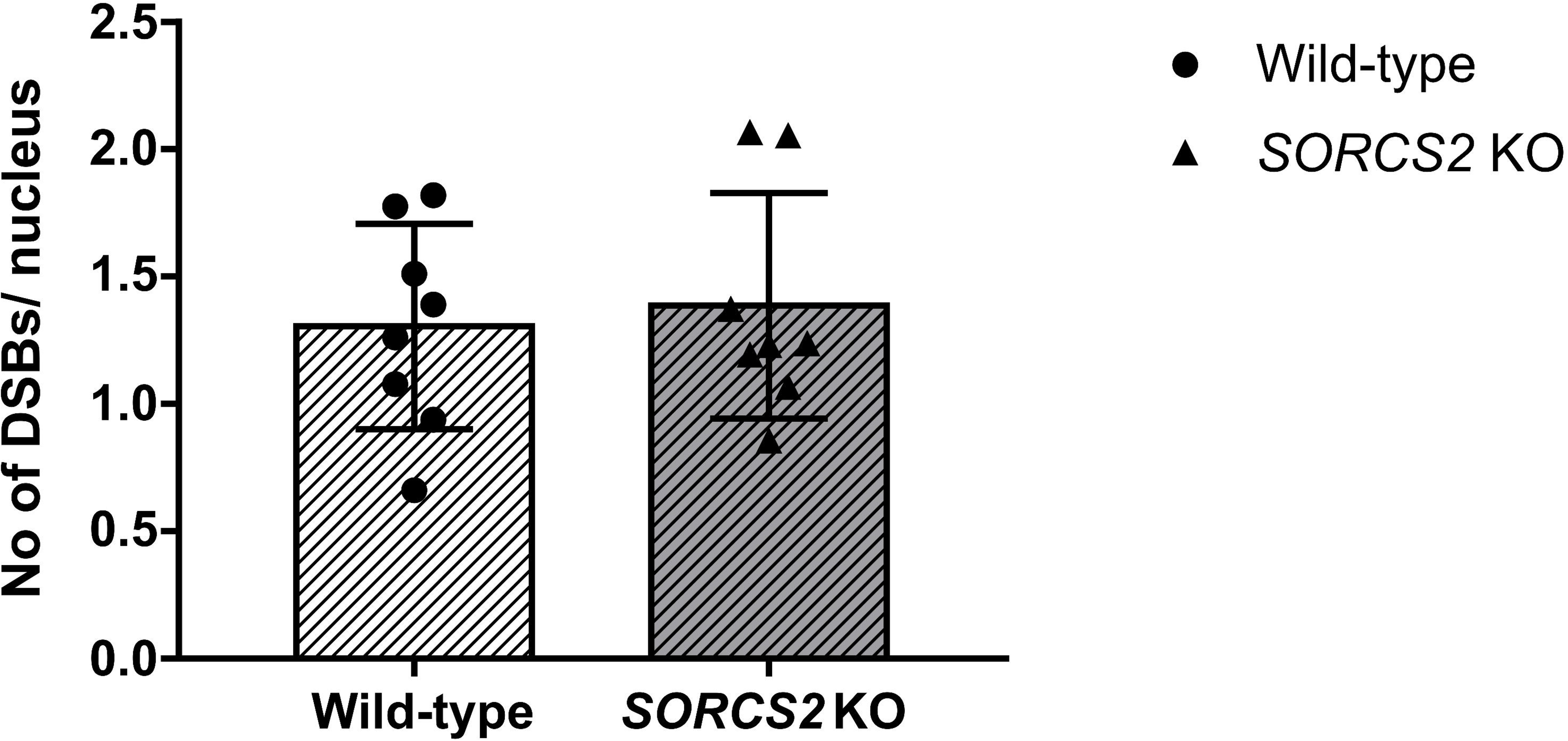
Treatment with Glycine has no differential effect on DNA DSB formation in *SORCS2* knock-out (KO) LUHMES neurons. No significant difference in the number of DSBs (γH2A.X/53BP1-positive foci) per nucleus was identified between wild-type (WT) (white bar) and *SORCS2* KO (grey bar) LUHMES neurons (day 14) following treatment with Glycine. Error bars represent means ± SD; n = 8 independent cell lines per genotype. Unpaired Student’s t-test, p > 0.05

Finally, given the potential negative impact of DSB formation on neuronal function and survival, we examined the effect of knocking out *SORCS2* on the overall neuronal viability both at early (day 6) and late (day 14) stages of differentiation. At day 6, there was no significant difference in the viability of wild-type neurons compared to that of the *SORCS2* knock-out clones (t_16_ = 0.296, p = 0.771; Fig. 5a). However, at day 14, we detected a significant reduction in the viability of *SORCS2*^-/-^ clones compared to controls (t_15_ = 3.387, p = 0.004; Fig. 5b).

**Fig. 5.**
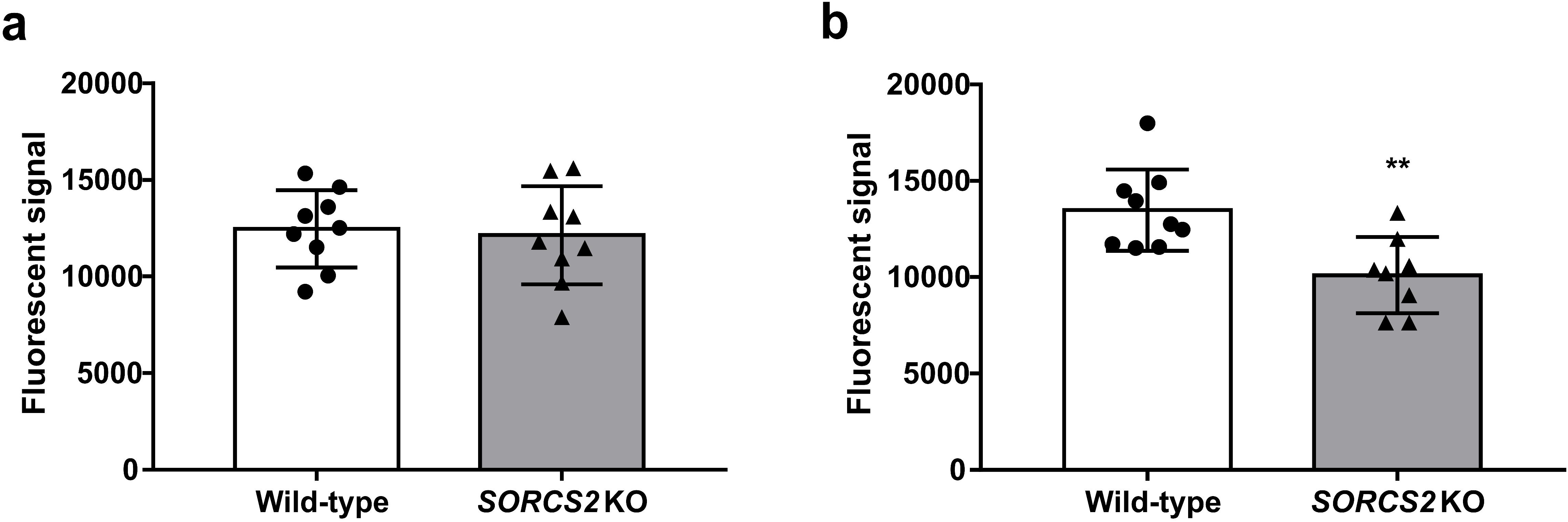
Knocking out *SORCS2* is associated with decreased neuronal viability at late (day 14), but not early (day 6) stages of neuronal differentiation. Neuronal viability of wild-type (WT) (white bar) and *SORCS2* knock-out (KO) (grey bar) LUHMES neurons measured at early (day 6) (a) and late (day 14) (b) stages of differentiation. ** p < 0.01 (unpaired Student’s t-test); Error bars represent means ± SD; n = 8-9 independent cell lines per genotype

## Discussion

Our data in wild-type mice are in agreement with the results of Suberbielle et al (2013), who showed that exploration of a novel environment is associated with the acquisition of DNA DSBs, which are repaired after a recovery period. We found, however, no evidence to support our initial hypothesis that *Sorcs2* knock-out mice would show a greater number of breaks associated with the exploratory behaviour or impaired recovery from this experience. In contrast, somewhat surprisingly, we observed higher levels of DNA DSBs in the DG of *Sorcs2*^*-/-*^ mice that remained in their home cage. We subsequently confirmed this in an independent set of knock-out and wild-type mice.

We next investigated whether higher levels of DNA DSBs would be also found in human neurons lacking *SORCS2*. DNA DSBs were rare in both mutant and wild-type lines, as has been reported previously for rat primary cortical neurons (Crowe et al 2006), and there was no detectable difference in γH2A.X immunoreactivity between the genotypes. As expected, treatment with the TopoIIβ inhibitor, etoposide, led to an increase in the number of breaks in both lines. The *SORCS2*^*-/-*^ lines, however, had significantly more breaks following etoposide treatment. Despite the increased number of TopoIIβ-dependent breaks in the knock-out cell lines, there was no difference in *TOP2B* expression levels in mutant lines either before or after treatment with etoposide. As enhanced TopoIIβ activity and DSB levels have been observed following stimulation of neuronal activity (Madabhushi et al 2015), we next investigated whether stimulation of neuronal activity would lead to a differential response in the neurons lacking SORCS2. We found no evidence that loss of SORCS2 rendered the human neurons more susceptible to neuronal activity-evoked DNA DSBs. This result is in keeping with our finding that *Sorcs2*^*-/-*^ mice appeared to have a similar number of DNA DSBs to wild-type mice following exploration of a novel environment; however, further work is required to determine both the impact of neuronal activation in mice and in other types of human neurons.

There are a number of potential explanations for the link between SORCS2 loss and DNA DSBs. Previous work (Malik et al 2019) implicated SorCS2 in protection against the oxidative stress-induced DNA damage and neuronal loss caused by a PTZ-induced kindling paradigm. Similarly, Smith *et al*. (2018) showed that *SORCS2* expression is stimulated by other stressors, such as alcohol and DEXA (Smith et al 2018). DEXA administration induces DNA damage, which can be prevented by application of reactive oxygen species (ROS) blockers (Ortega-Martínez 2015), thus SORCS2 loss may exacerbate the effect of cellular stressors on DNA damage. Previous work (Morotomi-Yano et al 2018) provides evidence for the participation of Topo IIβ in the cellular response to DSBs induced by laser microirradiation. It is possible, therefore, that etoposide treatment brings to light topoisomerase-mediated repair of breaks caused by loss of SORCS2, but independently of topoisomerase activity. Future experiments could test this hypothesis. Another possibility is that SORCS2 loss impacts the number of DNA DSBs through loss of interaction with DNA repair proteins. SORCS2 has been shown to co-localise with the transactivation response DNA-binding protein of 43kDa (TDP-43) in ALS post-mortem brains (Miki et al 2018). TDP-43 is an RNA/DNA-binding protein that has recently been implicated in DSB repair (Mitra et al 2019). SORCS2 also interacts with Heterogeneous Nuclear Ribonucleoprotein U (hnRNP-U) (Fasci et al 2018). This DNA and RNA binding protein interacts with NEIL1, a DNA glycosylase implicated in the repair of DNA damaged by reactive oxygen species, stimulating its base excision activity (Hegde et al 2012). Given the role of the VPS10P family in intracellular trafficking, future work could investigate whether SORCS2 is involved in trafficking the above proteins.

While the cellular mechanism underlying the increase in DNA DSBs associated with SORCS2 loss is still uncertain, it is of interest that mature (but not immature) *SORCS2*^*-/-*^ neurons showed decreased viability, in keeping with findings in mouse primary neurons lacking Sorcs2, which show higher rates of apoptosis (independent of autophagy) when subject to lysosomal stressors (Almeida at al., submitted). The maintenance of genome integrity is very important, particularly for post-mitotic long-lived cells, such as neurons, and DNA damage is linked to neurodegenerative disorders, ageing and decreased expression of genes important for brain maintenance and function (Madabhushi et al 2015). Further work is required, however, to investigate which aspect(s) of SORCS2 function underlie the observed decrease in viability.

This study is subject to a number of limitations. An important factor is the small number of replicates performed for the animal-based experiments, in particular. It is notable however that the set up was sufficient to reproduce the pattern seen by Suberbielle et al. in wild-type mice undergoing the novel environment task (Suberbielle et al 2013) and that we replicated the finding of increased numbers of breaks in the mutant mice that remained in the home cage in an independent set of mice. It is also notable that experiments performed in mice and a human cell line lacking SORCS2 both showed that SORCS2 loss was associated with a greater number of DNA DSBs. Further work is required to determine whether there are common or unique mechanisms underlying the findings.

In summary, we have shown that SorCS2 loss in mice leads to higher levels of γH2A.X-positive DNA breaks. Loss of SORCS2 in human neurons led to an increase in the number of TopoIIβ-dependent breaks and decreased neuronal viability. Our findings in both species suggest that the impact of SORCS2 loss is not mediated by a differing response to neuronal activation. An increase in DNA DSBs may lead to an altered transcriptional profile, affect genome integrity and ultimately lead to cell death. In agreement with this notion, DNA damage is increasingly being linked to cognitive impairment, dementia and other neurodegenerative disorders (Mullaart et al 1990; Adamec et al 1999; Madabhushi et al 2014; Shanbhag et al 2019; Thadathil et al 2019), and attenuating the DNA damage response to DSBs has been demonstrated to be protective in models of several neurodegenerative disorders (Tuxworth et al 2019). Our findings are in keeping with the known involvement of other sortilin family members in cognition, ageing and neurodegenerative disorders and with the recent finding that SNPs in *SORCS2* are involved in epistatic interactions associated with pathological hallmarks of Alzheimer’s disease (Wang et al 2020). Future experimental work should assess hypotheses based around SORCS2’s role in the cellular response to stress and/or DNA repair pathways and measure the impact of loss of Sorcs2 on the epigenome and transcriptome of the ageing dentate gyrus.

## Declarations

### Availability of data and materials

Please contact author for data requests.

### Code availability

Not applicable.

### Authors’ contributions

KOG and KLE conceived and planned the experiments. KOG performed the majority of the experiments and data analysis. SG provided the mice and DO and MK performed the behavioural experiments. SMA, JP, AP, PG and DP contributed to the execution of the experiments. RMW and MLB performed the statistical analysis of the mouse data. TSJ provided materials and support during assay optimisation. SG, CMA, TSJ and DJP contributed through strategic discussions. KOG and KLE wrote the manuscript with input from all authors.

### Ethics approval

All experiments were approved by the Danish Animal Experiments Inspectorate under the Ministry of Justice (Permits 2011/561-119, 2016-15-0201-01127 and 2017-15-0201-01192) and carried out according to the ARRIVE guidelines.

### Consent to participate

Not applicable.

### Consent for publication

Not applicable.

## Abbreviations

53BP1: p53-binding protein 1
ADHD: Attention deficit hyperactivity disorder
ALS: Amyotrophic lateral sclerosis
BDNF: Brain-derived neurotrophic factor
BSA: Bovine serum albumin
Ct: Cycle threshold
DEXA: Dexamethasone
DG: Dentate gyrus
DSB: Double-strand break
EV: Empty vector
FACS: Fluorescence-activated cell sorting
GDNF: Glial cell-derived neurotrophic factor
gRNA: Guide RNA
GWAS: Genome-wide association study
hnRNP-U: Heterogeneous nuclear ribonucleoprotein U
KCl: Potassium chloride
KO: Knock-out
LTD: Long-term depression
LTP: Long-term potentiation
LUHMES: Lund human mesencephalic
MSN: Medium spiny neurons
NHEJ: Non-homologous end joining
NMDAR: N-methyl-D-aspartate receptor
PBS: Phosphate-buffered saline
PLO: Poly-L-Ornithine
PSD: Post-synaptic density
PTZ: Pentylenetetrazol
ROS: Reactive oxygen species
SNP: Single-nucleotide polymorphism
TBS: Tris-Buffered Saline
TDP-43: Transactivation response DNA-binding protein of 43kDa
TH^+^: Tyrosine hydroxylase-positive
TopoIIβ: Topoisomerase IIβ
TrkB: Tropomyosin receptor kinase B
Vps10p: Vacuolar protein sorting (VPS) 10p
VTA: Ventral tegmental area
WT: Wild-type

## Acknowledgements

We thank Dr Ian Adams for helpful discussions regarding scientific strategy and the manuscript and Dr James Crichton for support with protocol optimisation and helpful comments on the manuscript. KOG received salary support from an Alzheimer’s Research UK award (ARUK-PPG2019B-015) to KLE. TSJ is funded by the UK Dementia Research Institute which receives its funding from DRI Ltd, funded by the UK Medical Research Council, Alzheimer’s Society, and Alzheimer’s Research UK, and the European Research Council (ERC) under the European Union’s Horizon 2020 research and innovation programme (Grant agreement No. 681181).

## References

Adamec E, Vonsattel JP, Nixon RA (1999) DNA strand breaks in Alzheimer’s disease. Brain Res 849:67–77. doi: 10.1016/s0006-8993(99)02004-1

Alemany S, Ribasés M, Vilor-Tejedor N, et al (2015) New suggestive genetic loci and biological pathways for attention function in adult attention-deficit/hyperactivity disorder. Am J Med Genet B, Neuropsychiatr Genet 168:459–470. doi: 10.1002/ajmg.b.32341

Crowe SL, Movsesyan VA, Jorgensen TJ, Kondratyev A (2006) Rapid phosphorylation of histone H2A.X following ionotropic glutamate receptor activation. Eur J Neurosci 23:2351–2361. doi: 10.1111/j.1460-9568.2006.04768.x

Davies G, Lam M, Harris SE, et al (2018) Study of 300,486 individuals identifies 148 independent genetic loci influencing general cognitive function. Nat Commun 9:2098. doi: 10.1038/s41467-018-04362-x

Deinhardt K, Kim T, Spellman DS, et al (2011) Neuronal growth cone retraction relies on proneurotrophin receptor signaling through Rac. Sci Signal 4:ra82. doi: 10.1126/scisignal.2002060

Duncan L, Yilmaz Z, Gaspar H, et al (2017) Significant Locus and Metabolic Genetic Correlations Revealed in Genome-Wide Association Study of Anorexia Nervosa. Am J Psychiatry 174:850–858. doi: 10.1176/appi.ajp.2017.16121402

Fabbri C, Serretti A (2016) Genetics of long-term treatment outcome in bipolar disorder. Prog Neuropsychopharmacol Biol Psychiatry 65:17–24. doi: 10.1016/j.pnpbp.2015.08.008

Fabbri C, Tansey KE, Perlis RH, et al (2018) New insights into the pharmacogenomics of antidepressant response from the GENDEP and STAR*D studies: rare variant analysis and high-density imputation. Pharmacogenomics J 18:413–421. doi: 10.1038/tpj.2017.44

Fasci D, van Ingen H, Scheltema RA, Heck AJR (2018) Histone interaction landscapes visualized by crosslinking mass spectrometry in intact cell nuclei. Mol Cell Proteomics 17:2018–2033. doi: 10.1074/mcp.RA118.000924

Firsanov DV, Solovjeva LV, Svetlova MP (2011) H2AX phosphorylation at the sites of DNA double-strand breaks in cultivated mammalian cells and tissues. Clin Epigenetics 2:283–297. doi: 10.1007/s13148-011-0044-4

Glerup S, Bolcho U, Mølgaard S, et al (2016) SorCS2 is required for BDNF-dependent plasticity in the hippocampus. Mol Psychiatry 21:1740–1751. doi: 10.1038/mp.2016.108

Glerup S, Nykjaer A, Vaegter CB (2014a) Sortilins in neurotrophic factor signaling. Handb Exp Pharmacol 220:165–189. doi: 10.1007/978-3-642-45106-5_7

Glerup S, Olsen D, Vaegter CB, et al (2014b) SorCS2 regulates dopaminergic wiring and is processed into an apoptotic two-chain receptor in peripheral glia. Neuron 82:1074–1087. doi: 10.1016/j.neuron.2014.04.022

Greenwood TA, Lazzeroni LC, Maihofer AX, et al (2019) Genome-wide Association of Endophenotypes for Schizophrenia From the Consortium on the Genetics of Schizophrenia (COGS) Study. JAMA Psychiatry 76:1274–1284. doi: 10.1001/jamapsychiatry.2019.2850

Hegde ML, Banerjee S, Hegde PM, et al (2012) Enhancement of NEIL1 protein-initiated oxidized DNA base excision repair by heterogeneous nuclear ribonucleoprotein U (hnRNP-U) via direct interaction. J Biol Chem 287:34202–34211. doi: 10.1074/jbc.M112.384032

Hermey G (2009) The Vps10p-domain receptor family. Cell Mol Life Sci 66:2677–2689. doi: 10.1007/s00018-009-0043-1

Karlsson Linnér R, Biroli P, Kong E, et al (2019) Genome-wide association analyses of risk tolerance and risky behaviors in over 1 million individuals identify hundreds of loci and shared genetic influences. Nat Genet 51:245–257. doi: 10.1038/s41588-018-0309-3

Lotharius J, Barg S, Wiekop P, et al (2002) Effect of mutant alpha-synuclein on dopamine homeostasis in a new human mesencephalic cell line. J Biol Chem 277:38884–38894. doi: 10.1074/jbc.M205518200

Ma Q, Yang J, Milner TA, et al (2017) SorCS2-mediated NR2A trafficking regulates motor deficits in Huntington’s disease. JCI Insight. doi: 10.1172/jci.insight.88995

Madabhushi R, Gao F, Pfenning AR, et al (2015) Activity-Induced DNA Breaks Govern the Expression of Neuronal Early-Response Genes. Cell 161:1592–1605. doi: 10.1016/j.cell.2015.05.032

Madabhushi R, Pan L, Tsai L-H (2014) DNA damage and its links to neurodegeneration. Neuron 83:266–282. doi: 10.1016/j.neuron.2014.06.034

Malik AR, Szydlowska K, Nizinska K, et al (2019) SorCS2 Controls Functional Expression of Amino Acid Transporter EAAT3 and Protects Neurons from Oxidative Stress and Epilepsy-Induced Pathology. Cell Rep 26:2792–2804.e6. doi: 10.1016/j.celrep.2019.02.027

Miki Y, Mori F, Seino Y, et al (2018) Colocalization of Bunina bodies and TDP-43 inclusions in a case of sporadic amyotrophic lateral sclerosis with Lewy body-like hyaline inclusions. Neuropathology 38:521–528. doi: 10.1111/neup.12484

Mitra J, Guerrero EN, Hegde PM, et al (2019) Motor neuron disease-associated loss of nuclear TDP-43 is linked to DNA double-strand break repair defects. Proc Natl Acad Sci USA 116:4696– 4705. doi: 10.1073/pnas.1818415116

Montecucco A, Zanetta F, Biamonti G (2015) Molecular mechanisms of etoposide. EXCLI J 14:95–108. doi: 10.17179/excli2015-561

Morotomi-Yano K, Saito S, Adachi N, Yano K-I (2018) Dynamic behavior of DNA topoisomerase IIβ in response to DNA double-strand breaks. Sci Rep 8:10344. doi: 10.1038/s41598-018-28690-6

Mullaart E, Boerrigter ME, Ravid R, et al (1990) Increased levels of DNA breaks in cerebral cortex of Alzheimer’s disease patients. Neurobiol Aging 11:169–173. doi: 10.1016/0197-4580(90)90542-8

Olsen D, Kaas M, Lundhede J, et al (2019) Reduced alcohol seeking and withdrawal symptoms in mice lacking the BDNF receptor sorcs2. Front Pharmacol 10:499. doi: 10.3389/fphar.2019.00499

Olsen D, Wellner N, Kaas M, et al (2021) Altered dopaminergic firing pattern and novelty response underlie ADHD-like behavior of SorCS2-deficient mice. Transl Psychiatry 11:74. doi: 10.1038/s41398-021-01199-9

Ortega-Martínez S (2015) Dexamethasone acts as a radiosensitizer in three astrocytoma cell lines via oxidative stress. Redox Biol 5:388–397. doi: 10.1016/j.redox.2015.06.006

Rampersad SN (2012) Multiple applications of Alamar Blue as an indicator of metabolic function and cellular health in cell viability bioassays. Sensors (Basel) 12:12347–12360. doi: 10.3390/s120912347

Scholz D, Pöltl D, Genewsky A, et al (2011) Rapid, complete and large-scale generation of post-mitotic neurons from the human LUHMES cell line. J Neurochem 119:957–971. doi: 10.1111/j.1471-4159.2011.07255.x

Shanbhag NM, Evans MD, Mao W, et al (2019) Early neuronal accumulation of DNA double strand breaks in Alzheimer’s disease. Acta Neuropathol Commun 7:77. doi: 10.1186/s40478-019-0723-5

Smith AH, Ovesen PL, Skeldal S, et al (2018) Risk locus identification ties alcohol withdrawal symptoms to SORCS2. Alcohol Clin Exp Res 42:2337–2348. doi: 10.1111/acer.13890

Suberbielle E, Sanchez PE, Kravitz AV, et al (2013) Physiologic brain activity causes DNA double-strand breaks in neurons, with exacerbation by amyloid-β. Nat Neurosci 16:613–621. doi: 10.1038/nn.3356

Thadathil N, Hori R, Xiao J, Khan MM (2019) DNA double-strand breaks: a potential therapeutic target for neurodegenerative diseases. Chromosome Res 27:345–364. doi: 10.1007/s10577-019-09617-x

Tuxworth RI, Taylor MJ, Martin Anduaga A, et al (2019) Attenuating the DNA damage response to double-strand breaks restores function in models of CNS neurodegeneration. Brain Commun 1:fcz005. doi: 10.1093/braincomms/fcz005

Wang H, Yang J, Schneider JA, et al (2020) Genome-wide interaction analysis of pathological hallmarks in Alzheimer’s disease. Neurobiol Aging 93:61–68. doi: 10.1016/j.neurobiolaging.2020.04.025

Yang J, Ma Q, Dincheva I, et al (2020) SorCS2 is required for social memory and trafficking of the NMDA receptor. Mol Psychiatry. doi: 10.1038/s41380-020-0650-7

