## Supplementary Material for "Loss of SORCS2 is associated with neuronal DNA double-strand breaks"

I Supplementary Figures and Figure Legends

**Suppl. Fig. 1**

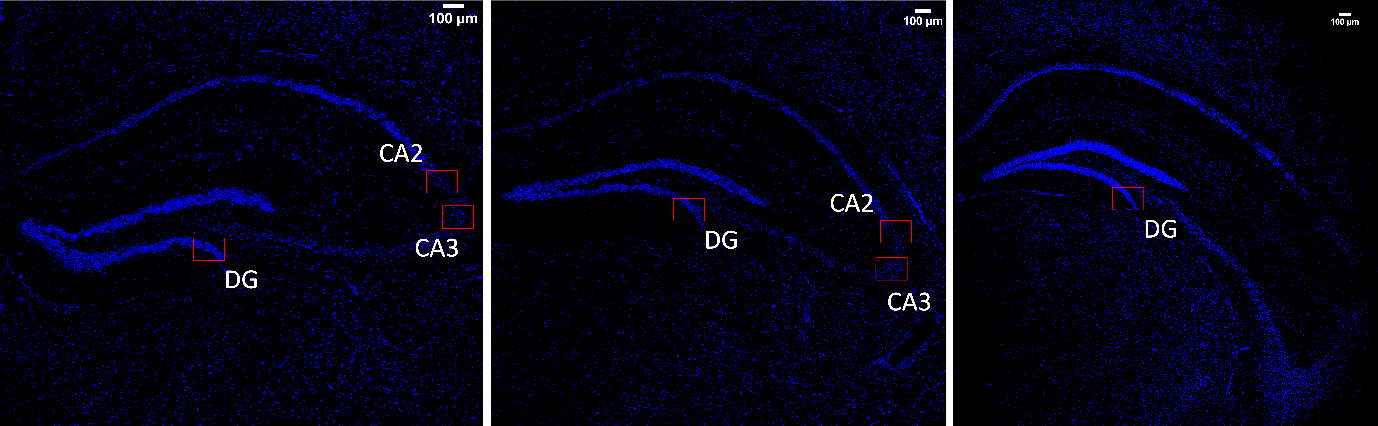

Overview images of the areas of the hippocampus immunostained for γH2A.X and counterstained with DAPI. Three (left to right) 14μm thick sections were obtained at intervals of 700μm, starting from the first section in which all regions of the hippocampus (i.e. CA1, CA2 and CA3) was visible (left section). The approximate location of the confocal images obtained within the DG, CA2 and CA3 regions is indicated with red boxes. 10x magnification; scale bars: 10μm

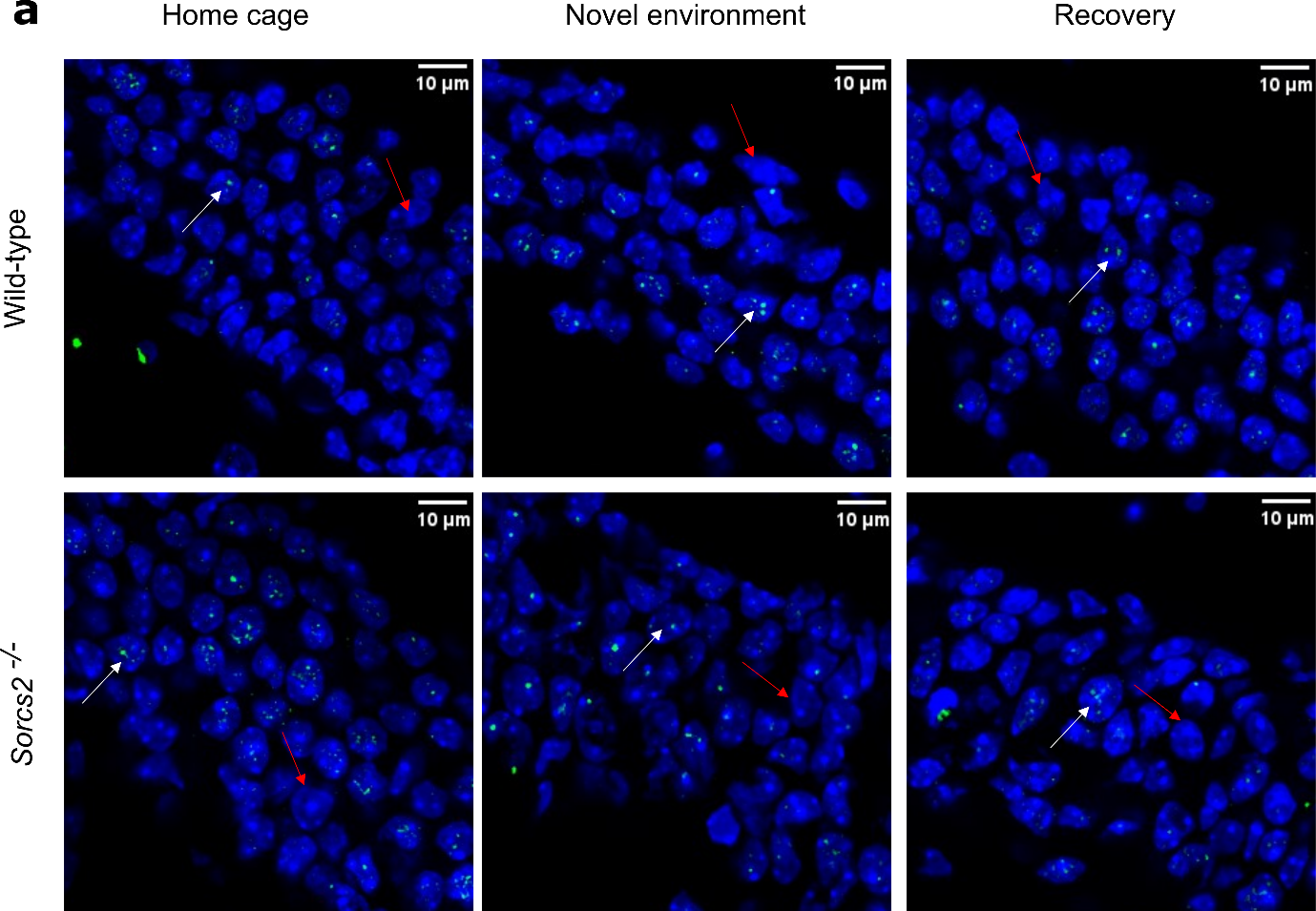

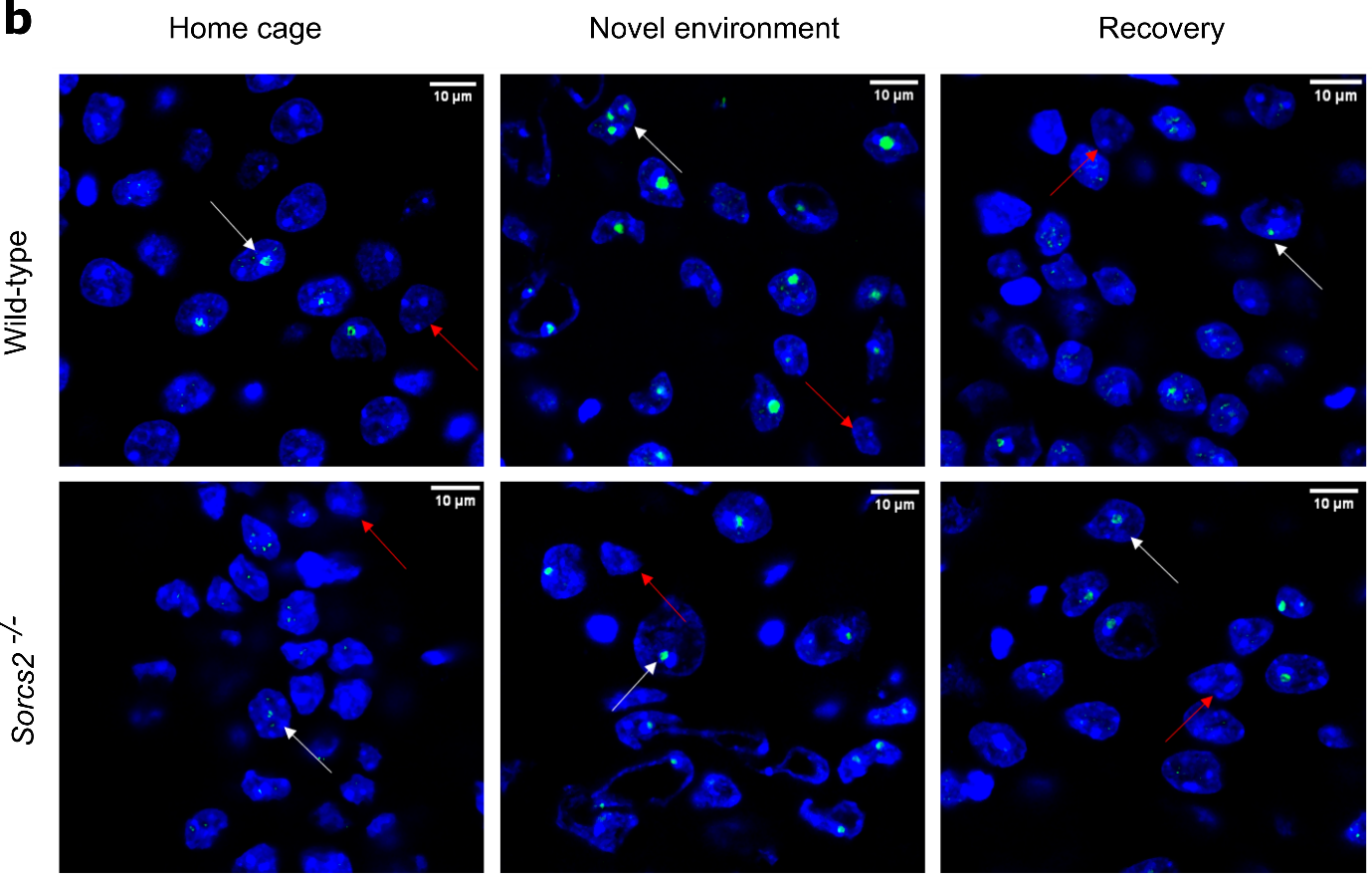

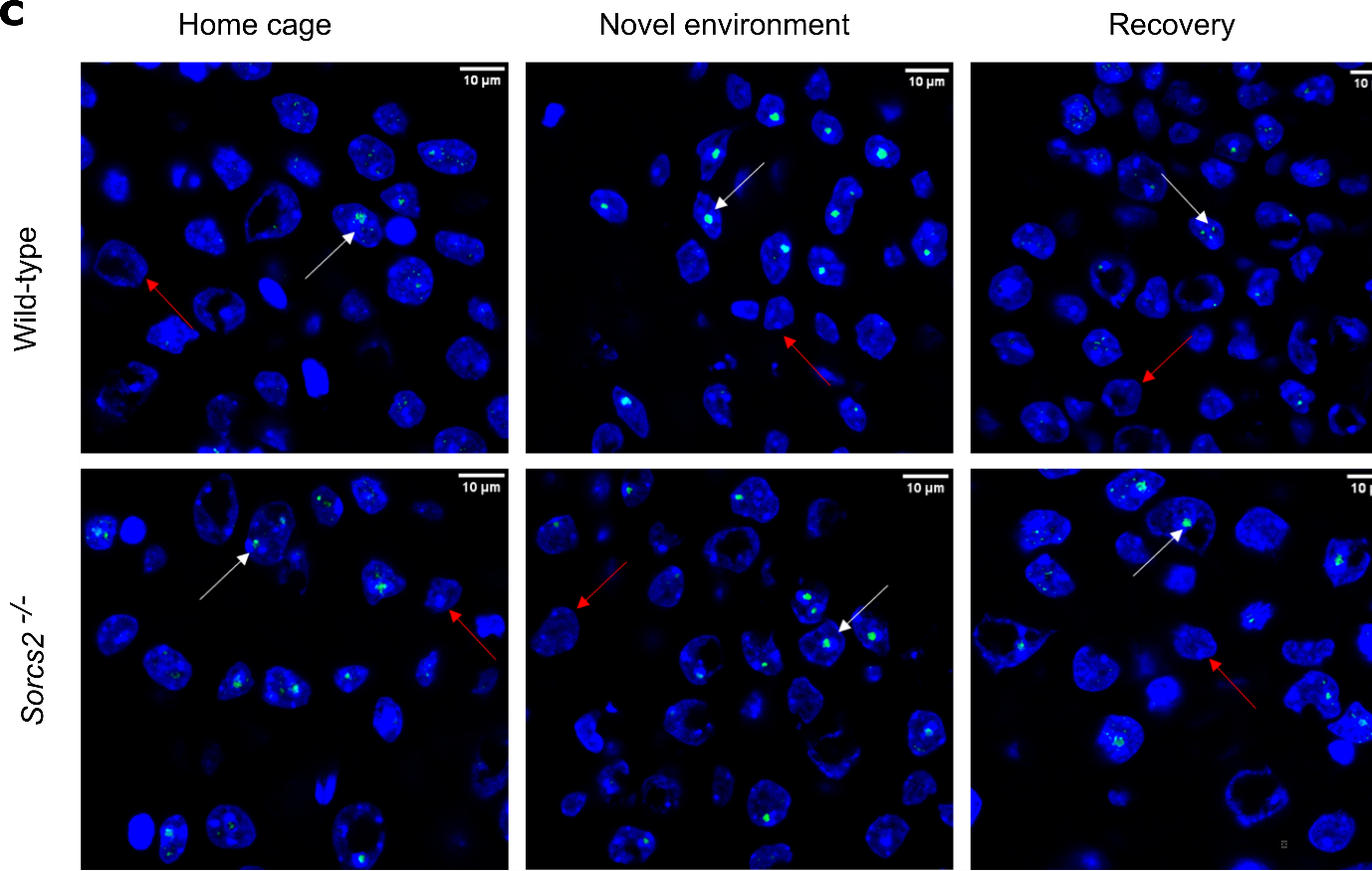

**Suppl. Fig. 2** DSB formation in wild-type (WT) and *Sorcs2*^-/-^ mice at baseline, following exploratory activity and recovery in the home cage. Single planes from the maximum projections of representative confocal images taken from the DG (a), CA2 (b) and CA3 (c) of WT (top row) and *Sorcs2*^-/-^ (bottom row) for mice from one of the three experimental conditions- home cage (left column), novel environment (middle column) and recovery (right column). γH2A.X-positive foci (green) were localised within nuclei (blue). When counting positive nuclei, due to the high nuclei density, each plane was examined individually. The nuclei positive for γH2A.X in each plane were marked while counting to avoid oversampling. White and red arrows point towards γH2A.X-positive and negative nuclei, respectively. 60x magnification; scale bars: 10μm

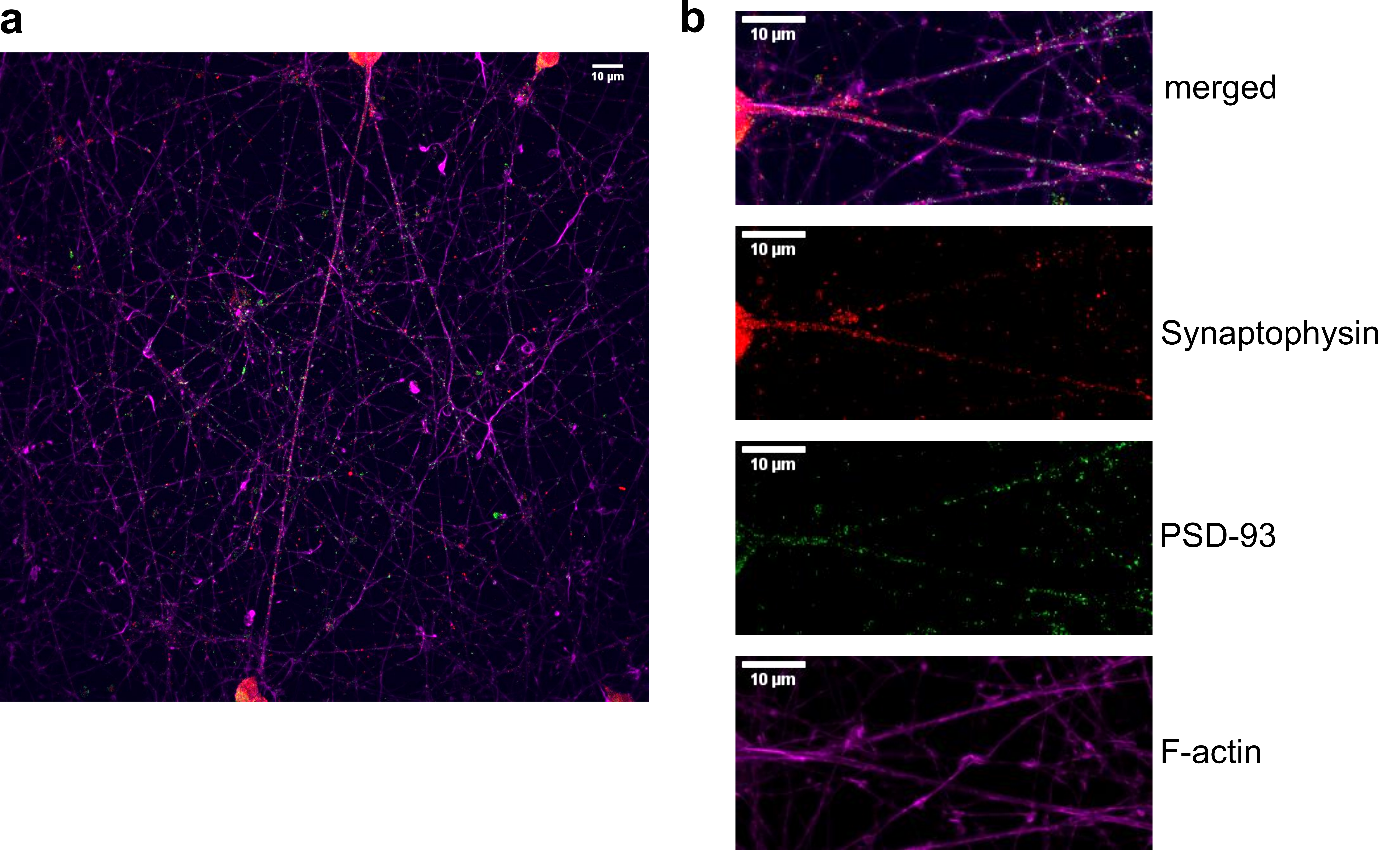

**Suppl. Fig. 3** Wild-type (WT) LUHMES neurons (day 14) stained for F-actin (magenta), PSD-93 (green), Synaptophysin (red), and DAPI (blue). An overview confocal image (a) and cropped images focusing on the axons (b). 60x magnification; scale bars: 10μm

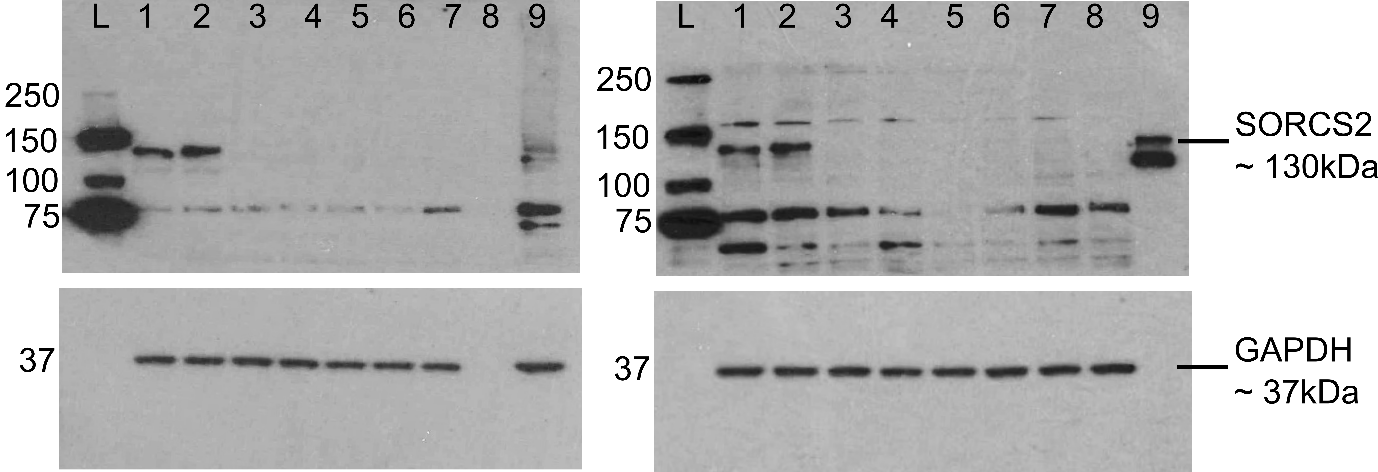

**Suppl. Fig. 4** Full-size western blots corresponding to the blots shown in Fig.2b and showing a complete loss of SORCS2 in the knock-out (KO) clones after targeting exon 1 (left) or exon 3 (right). Samples loaded on the blot on the left correspond to: 1 and 2 lysates obtained from wild-type (WT) LUHMES neurons (day 14), 3- 7- from *SORCS2* KO exon 1 clones 1- 5 (day 14) generated by targeting exon 1, 8- empty, 9- lysates obtained from iPSC-derived neurons. Samples loaded on the blot on the right correspond to: 1 and 2 lysates obtained from WT LUHMES neurons (day 14), samples 3- 8- from *SORCS2* KO exon 3 clones 1- 6 (day 14) generated by targeting exon 3. Sample 9 constitutes a positive control (protein lysate from HEK293 cells overexpressing *SORCS2*). ‘L’ stands for ladder in both blots. *SORCS2* exon 1 clone 4 and *SORCS2* exon 3 clone 5 did not survive neuronal differentiation and were not included in any subsequent experiments.

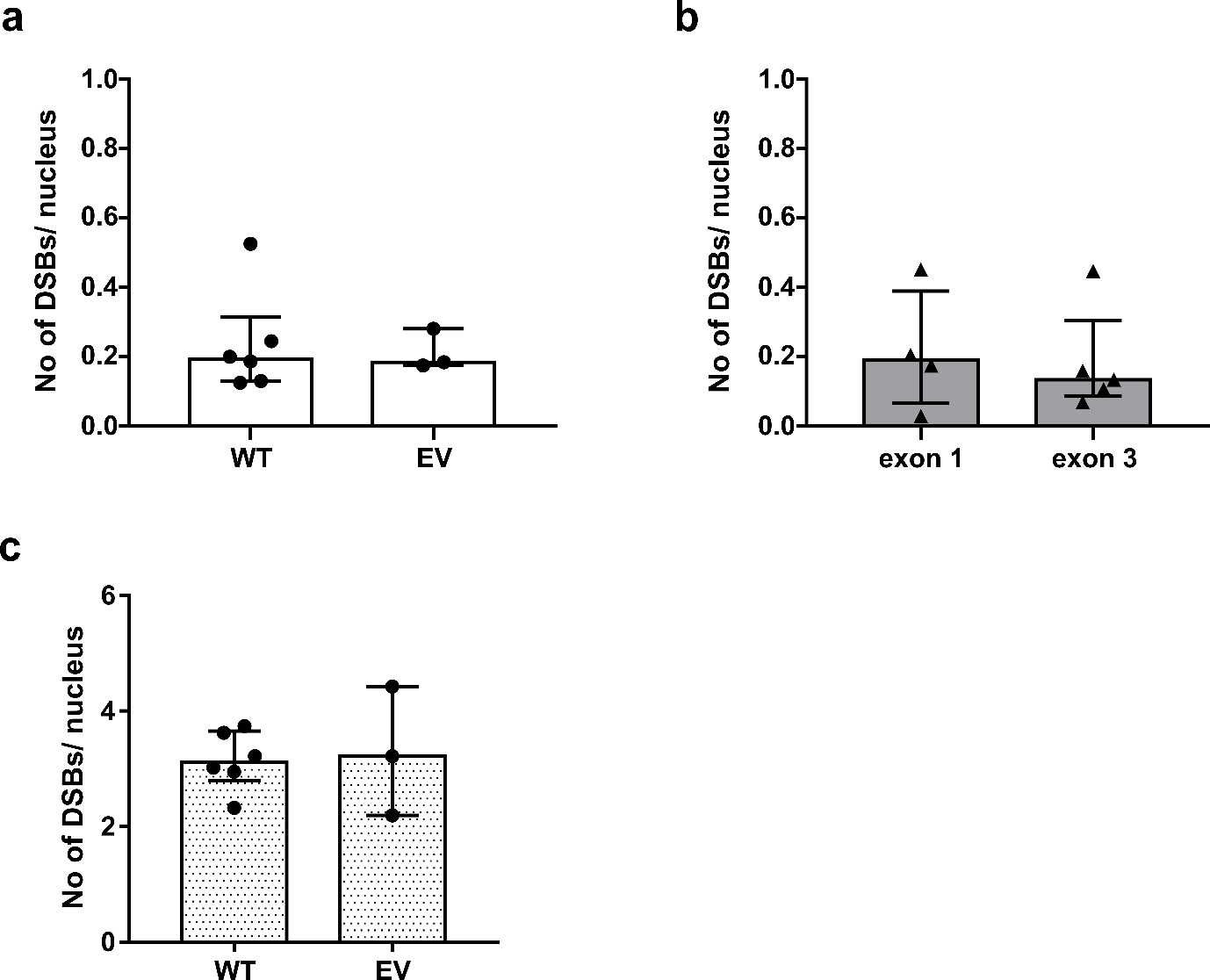

**Suppl. Fig. 5** DSB formation in wild-type (WT) and *SORCS2* knock-out (KO) LUHMES neurons. Number of DSBs (γH2A.X/53BP1-positive foci) per nucleus in untreated (a) WT (n= 6 independent cell lines) and empty vector (EV) controls (n= 3 independent cell lines) and (b) *SORCS2* KO LUHMES neurons (day 14) generated by targeting exon 1 (n=4 independent cell lines) or exon 3 (n=5 independent cell lines). (c) Number of DSBs (γH2A.X/53BP1-positive foci) per nucleus in etoposide-treated WT (n= 6 independent cell lines) and EV controls (n= 3 independent cell lines) LUHMES neuron day 14. Mann-Whitney test, p > 0.05. Approximately 100 nuclei counted per cell line. Error bars represent median with interquartile range

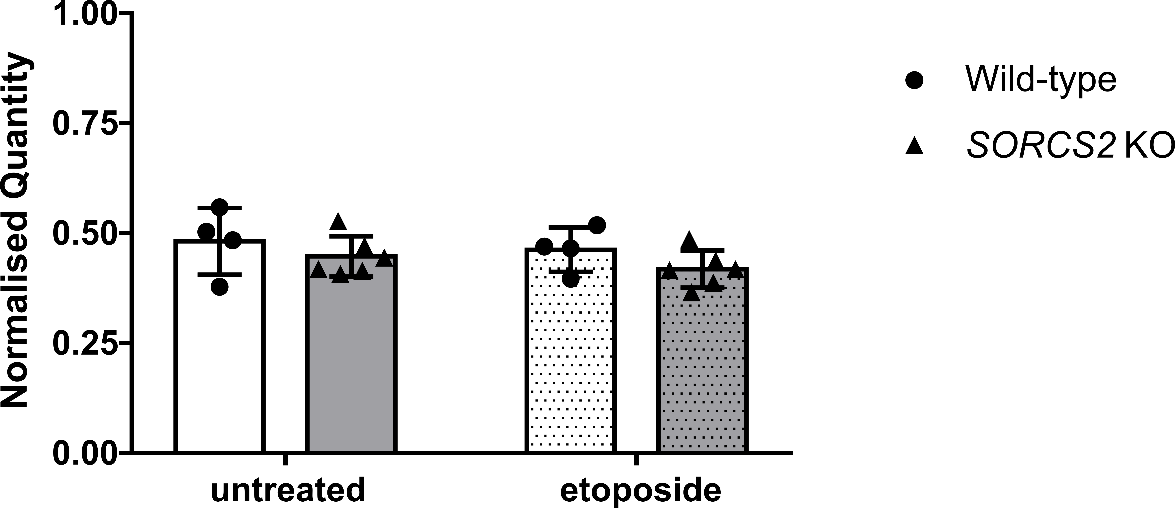

**Suppl. Fig. 6** Knocking out *SORCS2* has no effect on *TOP2B* expression levels. *TOP2B* expression levels in untreated (open bars) and etoposide-treated (dotted bars) WT and *SORCS2* KO LUHMES neurons (day 14). *TOP2B* expression was normalised to the expression of two reference genes- *SDHA* and *UBE4A*; No significant effect of the treatment (F_1,16_ = 0.978, p = 0.337), the genotype (F_1,16_  = 2.652, p = 0.123) or the interaction between the two (F_1,16_ = 0.043, p= 0.839) was identified (Two-way ANOVA). N= 4-6 independent WT and *SORCS2* KO lines, respectively; error bars represent means ± SD

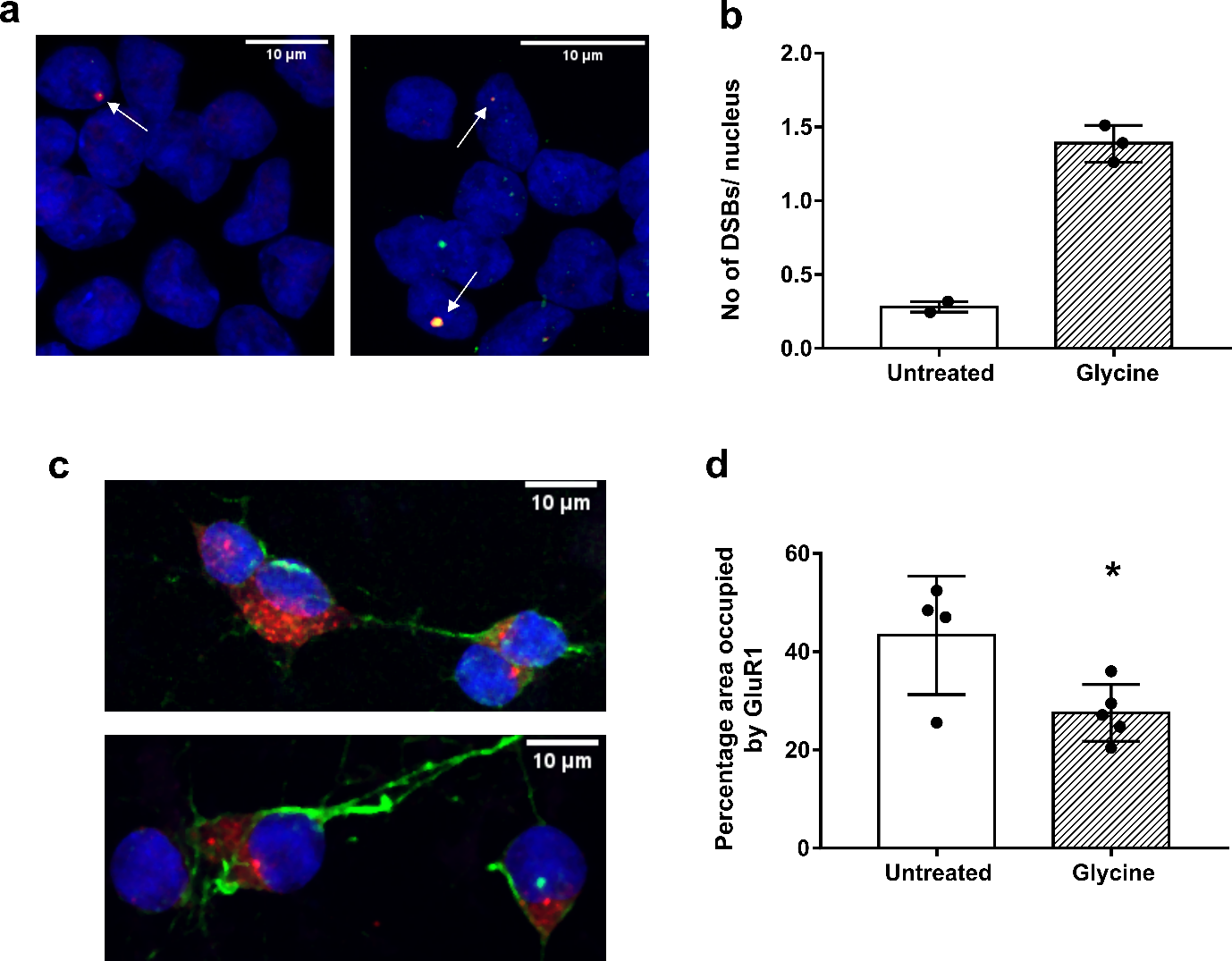

**Suppl. Fig. 7** Glycine treatment is associated with increased DSB formation and reduced somatic surface area occupied by AMPA receptors (GluR1) in wild-type (WT) LUHMES neurons (day 14). (a) Representative confocal images from untreated (left) and glycine-treated (right) WT LUHMES neurons (day 14) immunostained with γH2A.X (green) and 53BP1 (red), and counterstained with DAPI (blue). White arrows point towards γH2A.X/53BP1 dual positive. Images were taken at 100x magnification; scale bars: 10 μm. (b) Number of DSBs (γH2A.X/53BP1-positive foci) per nucleus in untreated and Glycine treated WT LUHMES neurons (day 14). Cells were treated with: Glycine (300µM) in Mg^2+^ free ACSF for 5min, followed by 15min recovery in ACSF with Mg^2+^. N = 2-3 independent WT lines, approximately 100 nuclei counted per cell line. Error bars represent median with interquartile range. (c) Representative confocal images from untreated (top) and glycine-treated (bottom) WT LUHMES neurons (day 14) immunostained with GluR1 (red) and F-actin (green), and counterstained with DAPI (blue). Images were taken at 100x magnification; scale bars: 10 μm. (d) Somatic surface area occupied by AMPA receptors (GluR1) in untreated and glycine-treated control (WT and empty vector [EV]) LUHMES neurons (day 14). * p< 0.05 (unpaired Student’s t-test); Error bars represent means ± SD

II Supplementary Tables

**Suppl. Table 1** Results of the tests for normality and variance heterogeneity and statistical analysis performed for each experiment (listed by figure number)

| **Figure number** | **Normal Distribution?** | **Homogeneity of variance?** | **Type of Statistical Analysis** |
| --- | --- | --- | --- |
| **Figure 1B**  n=3 | Not applicable- n too small | Not applicable- n too small | No statistical analysis |
| **Figure 1C**  WT mice (n=7)  KO mice (n=5) | Yes  Yes | No | Non-parametric |
| **Figure 3B**  Untreated WT (n=9)  Untreated KO (n=9) | No  Yes | Yes | Non-parametric |
| **Figure 3C**  Etoposide-treated WT (n=9)  Etoposide-treated KO (n=9) | Yes  Yes | Yes | Parametric |
| **Figure 3D**  Etoposide-treated KO exon 1 (n=4)  Etoposide-treated KO exon 3 (n=5) | Yes  No | Yes | Non-parametric |
| **Figure 4**  Gly- treated WT (n=8)  Gly- treated KO (n=8) | Yes  Yes | Yes | Parametric |
| **Figure 5A**  D6 WT (n=9)  D6 KO (n=9) | Yes  Yes | Yes | Parametric |
| **Figure 5B**  D14 WT (n=9)  D14 KO (n=8) | Yes  Yes | Yes | Parametric |
| **Suppl. Fig. 5A**  Untreated WT (n=6)  Untreated EV (n=3) | Not applicable- n too small | Not applicable- n too small | No statistical analysis |
| **Suppl. Fig. 5B**  Untreated exon 1 (n=4)  Untreated exon 3  (n=5) | Yes  No | No | Non-parametric |
| **Suppl. Fig. 5C**  Etoposide-treated WT (n=6)  Etoposide-treated EV  (n=3) | Not applicable- n too small | Not applicable- n too small | No statistical analysis |
| **Suppl. Fig. 6**  Untreated WT (n=4)  Etoposide-treated WT (n=4)  Untreated KO (n=6)  Etoposide-treated KO (n=6) | Yes  Yes  Yes  Yes | Yes | Parametric |
| **Suppl. Fig. 7 B**  n=2-3 | Not applicable- n too small | Not applicable- n too small | No statistical analysis |
| **Suppl. Fig. 7 D**  n=4-5 | Yes | Yes | Parametric |

**Suppl. Table 2** median and interquartile range (IQR) of the percentage γH2A.X-positive neurons in the DG, the CA2 and CA3 region of the hippocampus of wild-type and *Sorcs2*^-/-^ mice belonging to one of the three experimental groups- home cage, novel environment and recovery

| **Condition** | **Dentate gyrus** | | **CA2** | | **CA3** | |
| --- | --- | --- | --- | --- | --- | --- |
|  | **Median %** | **IQR** | **Median %** | **IQR** | **Median %** | **IQR** |
| **Wild-type mice** | | | | | | |
| **Home cage** | 31.503 | 4.210 | 38.472 | 6.356 | 38.472 | 6.356 |
| **Novel environment** | 59.427 | 2.325 | 57.033 | 4.220 | 57.033 | 4.220 |
| **Recovery** | 42.683 | 2.185 | 39.301 | 13.000 | 39.301 | 13.000 |
| ***Sorcs2*^-/-^ mice** | | | | | | |
| **Home cage** | 46.045 | 4.085 | 42.416 | 11.134 | 42.416 | 11.134 |
| **Novel environment** | 52.367 | 6.018 | 54.084 | 3.002 | 54.084 | 3.002 |
| **Recovery** | 45.603 | 1.051 | 40.809 | 6.026 | 40.809 | 6.026 |
